# A2Holo: Dual-Stream Flow Matching of 3D Structure and a Protein Language Model for Conformational Transition Prediction

**DOI:** 10.64898/2026.11.20.768033

**Authors:** Dongwoo Kim, Haelyn Kim, Jaemin Sim, Minjong Kim, Myunghyun Jang, Juyong Lee

## Abstract

Modelling a ligand-bound holo conformation from a ligand-free apo conformation is essential in structure-based drug discovery. We present A2Holo, which casts apo-to-holo prediction as flow matching between two experimental structures rather than generation from noise. It transplants the dual-stream joint attention of in-context image generation to residue-aligned protein language model embeddings and structure. Without ligand conditioning or stochastic sampling, A2Holo’s single deterministic trajectory reaches a median C*α* RMSD of 0.809 Å on the D3PM benchmark, the lowest among existing methods. Its TM-score exceeds even the best of 10 samples from every stochastic baseline on at least 144 of 150 pairs. Beyond performance, we directly measure the transplanted multimodal mechanism itself. Because sequence and structure are residue-aligned, its joint attention can be measured at residue resolution and tested for reproducibility across independently trained model variants. Within the structure stream, the attention a residue receives recovers moving residues better than the attention it emits, and this asymmetry is learned. A2Holo demonstrates a successful multimodal transplant from in-context image generation to protein conformational change, with strong deterministic apo-to-holo prediction. It also provides a quantitative account of how that mechanism organises residue-aligned sequence and structure around the residues that move.

## 1 Introduction

A protein that binds a ligand does not keep its shape. Under conformational selection followed by induced fit, the ligand selects from a pre-existing ensemble and binding then redistributes it (Boehr et al., 2009). One deposited unbound structure therefore does not fix the bound form on its own. Structure-based drug design needs that bound form, while the structure at hand is often the unbound one. What a ligand-free predictor must produce is the change between the two states (Figure 1). The sequence is shared by both states, and the apo coordinates fix only where the change starts, so the two inputs stand in different relations to it.

**Figure 1:**
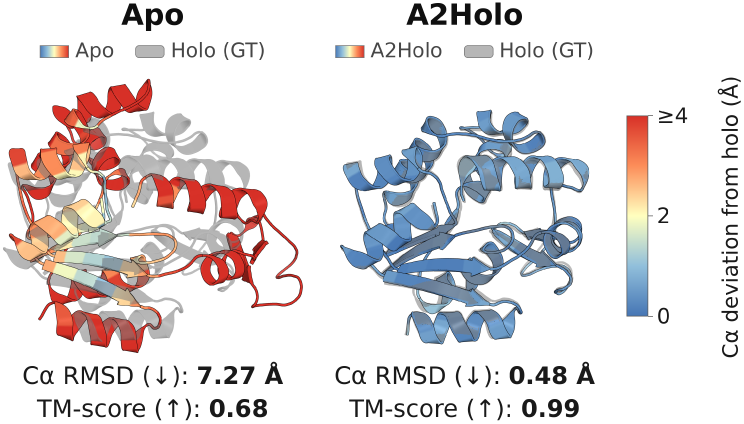
What the task asks for, on one pair, PDB 4AKE (apo) to 4X8L (holo), chain A in both. Grey is the bound (holo) deposition, the target. Colour is how far each residue sits from it. **Left**, the unbound (apo) deposition, the only structure the model is given. No ligand is given or drawn. Both are experimental. **Right**, A2Holo’s prediction from that input. The pair is chosen for legibility and is not typical. It moves 7.27 Å where the median test pair moves 4.46 Å, and the two numbers under each panel are this pair’s, not the benchmark’s, whose median is 0.809 Å (Section 3).

Neither experiment nor simulation closes the gap. Few proteins have both states deposited. The apo–holo benchmark used here holds 1591 pairs, curated by Somnath et al. (2023) from the D3PM database (Peng et al., 2022). Molecular dynamics reaches the transition only at prohibitive cost (Seeliger & de Groot, 2010). Surrogates (Lewis et al., 2025) return a distribution from the sequence alone, and picking the bound state out of it needs the answer.

Generative models pose the bound state as a stochastic sampling problem and, at their most accurate, condition on a ligand not yet known. The nearest exception in framework is Sesame (Miñán et al., 2025), a conditional flow-matching model on backbone frames that calls itself ligand-agnostic, though a ligand-defined pocket mask enters its preprocessing and it reports no sample count.

The two inputs are not redundant. A protein language model over the sequence already carries much of the apo structure, reaching a canonical correlation of 0.854 with frame-invariant apo geometry. It reaches 0.995 with the physicochemical block often concatenated beside it. Even so, neither input stands in for the other.

A model-free linear probe predicts the per-residue displacement profile with out-of-sample *R*^2^ of 0.196 from geometry and 0.137 from the language model, and both together reach 0.244 (Figure A1). Therefore, ligand-free apo-to-holo prediction can draw on both modalities. Protein models already use both. DPLM-2 (Wang et al., 2025) is one multi-modal diffusion language model over sequence and structure, and PT-DiT (Lin et al., 2025) codes both into one token per residue for a single-stream diffusion transformer.

We introduce A2Holo, which maps an apo structure to its holo form. Its sequence and structure streams meet in the dual-stream joint attention of in-context image generation (Esser et al., 2024; Black Forest Labs et al., 2025).

Whether that layout does in proteins what it does with text and pixels is a separate question, and this paper answers it by measurement. The two streams here run over the same residue index, so each quadrant of the joint map is attributable to one residue and the interaction can be measured per residue (Figure 4). We measure it on two axes, what the streams do to each other and what the stack does with it (Section 4). Both are measured on independently trained models (9 trainings over 5 depths).

We make one modelling contribution and two measured ones.

- A2Holo is the state of the art on the D3PM benchmark under the split used by prior work, and carries protein multimodality as two streams. In its double-stream blocks they keep independent weights and meet only inside joint attention. One deterministic trajectory, given no ligand, reaches the lowest median C*α* RMSD, 0.809 Å against 4.46 Å for the unchanged apo input. Its TM-score beats every stochastic baseline at its target-selected best of 10 samples on at least 144 of 150 pairs (Section 3, Table 1, Figures 3 and A3).
- The attention a residue pays and the attention paid to it are different quantities, which the usual row-plus-column sum collapses. The split is absent at random initialisation and appears once the model is trained (9 of 9 trainings). Within the structure stream the attention paid to a residue picks out the moving residues, overlapping them at 1.88–2.54 times the rate a random selection would (Figures 5 and 6), and in every training it exceeds every quantity indexed by a sequence token (9 of 9). Holding distance from the apo centroid fixed, received attention still tracks the change (9 of 9 trainings). What the sequence stream pays to the structure stream picks out the moving residues better than what it pays to itself (9 of 9). Attention marks *where* the structure changes, not *how* (Sections 4.2–4.4). All of it holds when the model is given the apo structure alone (Appendix A.18).
- The structure stream carries more about the coming change than the language model alone, with relative decodability below one for displacement and for total distance change alike (9 of 9 trainings). These effects reproduce across retrainings while their depth does not, with the received-attention peak anywhere from 0.333 to 0.909 of the double-stream stack and the decodability minimum from 0.348 to 0.957 of both stacks (Sections 4.1 and 4.5).

**Table 1:** Apo-to-holo C*α* prediction on the D3PM benchmark (150 test pairs). Baseline C*α* RMSDs are quoted from Nobis et al. (2025), Table 1 (Appendix A.8). *<*2 and *<*5 give the percentage of pairs under 2 and 5 Å (for SBAlign, of its pooled samples). A2Holo is the released checkpoint, seed 1, and its 5 seeds span medians 0.761–0.836 Å (Table A4). NFE counts network function evaluations.

|  | Cα RMSD (Å) |  |  |  |  |
|---|---|---|---|---|---|
| Method | Median | Mean | <2 | <5 | sampling |
| Apo input (no prediction) | 4.46 | 6.74 | 0% | 59% | — |
| EGNN | 19.99 | 21.37 | 1% | 1% | single |
| SBAlign | 3.8 | 4.98 | 0% | 69% | 10 samples, pooled |
| Sesame | 2.87 | 3.65 | 38% | 82% | not reported |
| ABM | 2.4 | 3.49 | 43% | 84% | single, 5 trainings |
| FDBM | 2.12 | 3.34 | 48% | 86% | single, 5 trainings |
| **A2Holo (ours)** | **0.809** | **2.357** | **66.7%** | **88.0%** | **deterministic, NFE 10** |

## 2 Related work

### One structure per target

Single-structure predictors return one conformer (Jumper et al., 2021; Abramson et al., 2024). That conformer is holo-biased on roughly two-thirds of targets (Saldaño et al., 2022). Docking takes the receptor conformation as an input (Morehead et al., 2026b). MSA subsampling surfaces alternatives and ranks them (del Alamo et al., 2022), though not by which one a ligand selects.

### Generative models and ensembles

Generative models take that up by sampling stochastically (Somnath et al., 2023; Nobis et al., 2025), conditioning on the ligand (Lu et al., 2024; Qiao et al., 2024; Corso et al., 2025), or emulating conformational ensembles from sequence (Jing et al., 2024; Lewis et al., 2025). Table 1 compares against two of these, the aligned Schrödinger bridge (SBAlign, Somnath et al., 2023) and the fractional diffusion bridge (FDBM, Nobis et al., 2025), and against an EGNN (Garcia Satorras et al., 2021) and augmented bridge matching (ABM, De Bortoli et al., 2023), which Somnath et al. (2023) and Nobis et al. (2025) respectively trained on the same split. Ensemble models cover the range of states rather than the transition (Wang et al., 2026a). Adjacent lines design new proteins (Jendrusch & Korbel, 2025; Zhang et al., 2025), predict one structure (Wang et al., 2026b), study sampling budgets (Liu et al., 2026) and survey flow matching (Morehead et al., 2026a). Two further apo-to-holo models remain. ApolloDiff withholds the ligand, as A2Holo does, but samples several structures per pair (Zhang et al., 2023). Sesame (Miñán et al., 2025), described in Section 1, is quoted in Table 1 from its published RMSDs. These works set out the prediction problem. The mechanism this paper studies comes from a different lineage.

### Joint attention and non-equivariance

Carrying two streams in one attention is an image-generation design. It runs from the diffusion transformer (Peebles & Xie, 2023), through one attention map over text and image tokens (Esser et al., 2024), to a double-then-single stack (Black Forest Labs et al., 2025). In those models joint attention is how text steers pixels. The block layout is adopted here, and Section 4 asks whether it does the same job. Shin et al. (2025) split that joint map into its four blocks, text and image as query and as key, and use them to edit images. An image model’s two streams are not in correspondence, a fixed block of encoder tokens against a grid of latents. Both streams here run over the same residue index, so a quadrant of the joint map is readable per residue, unlike a single-stream model that merges the two embeddings of each residue into one token (Jing et al., 2025). The other architectural choice, non-equivariance, is likewise prior art (Geffner et al., 2025; Jing et al., 2025), following Abramson et al. (2024).

### Reading attention as evidence

Reading that map as evidence is argued both ways (Jain & Wallace, 2019; Wiegreffe & Pinter, 2019), and probing has its own confounds (Belinkov, 2022). The closest precedent in proteins reads a language model’s attention by depth, and resolves depth on one checkpoint (Vig et al., 2021). What decides whether such a reading survives is the baseline it is measured against, and Belinkov (2022) names the one usually missing, several architectures trained on the same data. Every claim this paper makes is stated in a form that holds across independently trained models (9 trainings over 5 depths, Section 4.1).

## 3 Model and task

### Transport between two experimental structures

We pose apo-to-holo prediction as transport between two experimentally determined structures rather than as generation from noise (Appendix A.1). Each training pair supplies the apo and the holo coordinates of the same residues, in one shared frame (Appendix A.5). The model learns a velocity field **v***_θ_*(**x***_t_, t*) along the straight path that joins them. The interpolant carries no noise term, **x***_t_* = (1 − *t*) **x**_apo_ + *t* **x**_holo_. Its target velocity is the displacement **u***_t_* = **x**_holo_ − **x**_apo_. Supervision is applied in position space through **x**^_holo_ = **x**_apo_ + **v***_θ_*(**x***_t_, t*), so the field is trained to the constant velocity of the linear path (Lipman et al., 2023), with data at both endpoints (Liu et al., 2023; Albergo et al., 2025). We write the perresidue displacement as Δ**x***_i_* = **x**_holo*,i*_ − **x**_apo*,i*_. Time runs from the apo state at *t* = 0 to the holo state at *t* = 1, the reverse of SD3, from which the backbone is adapted (Esser et al., 2024). Inference integrates the field forward from **x**^(0)^ = **x**_apo_ in *K* = 10 equal forward-Euler steps,

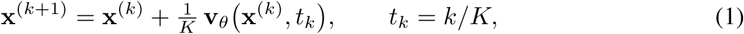

with no noise, ensembling, or sample selection (Appendix A.7 compares one and a hundred steps). Section 4 relies on the determinism, measuring a perturbation against a single unperturbed reference with no sampling variance to separate from the effect.

### Two streams, one attention over both

Two per-residue streams enter the network. The structure stream carries the current C*α* and C*β* coordinates under a Fourier feature encoding (Tancik et al., 2020), concatenated with per-residue node features. The sequence stream carries frozen 1536-dimensional per-residue representations from the Ankh3-large protein language model (Alsamkary et al., 2025), and both are projected to width 768. The flow time and a global token form one conditioning vector, which sets each block’s AdaLN shift, scale and gate. The global token is the Ankh3 output at its `[NLU]` position (Appendices A.2 and A.3). The network stacks 12 double-stream blocks followed by 12 single-stream blocks, with 8 attention heads and an MLP ratio of 4.0. Its layout is adopted from the FLUX.1 and SD3 image backbones (Black Forest Labs et al., 2025; Esser et al., 2024). A double-stream block keeps independent weights for the two streams and couples them only inside attention (Figure 2). Queries, keys and values are computed per stream, concatenated along the token axis, attended over once and split back. *The sequence stream is placed first in that concatenation*. Attention is computed one protein at a time, so each map spans 2*N* tokens for a protein of *N* residues and a residue attends only within its own chain. Every quadrant read in Section 4 is defined against this token order. Rotary position embeddings (Su et al., 2024) are one-dimensional over residue index and duplicated across the two streams, so residue *i* carries the same phase in both halves. The single-stream blocks act on the merged sequence. They compute attention and the feed-forward branch in parallel from the same modulated input (Dehghani et al., 2023).

**Figure 2:**
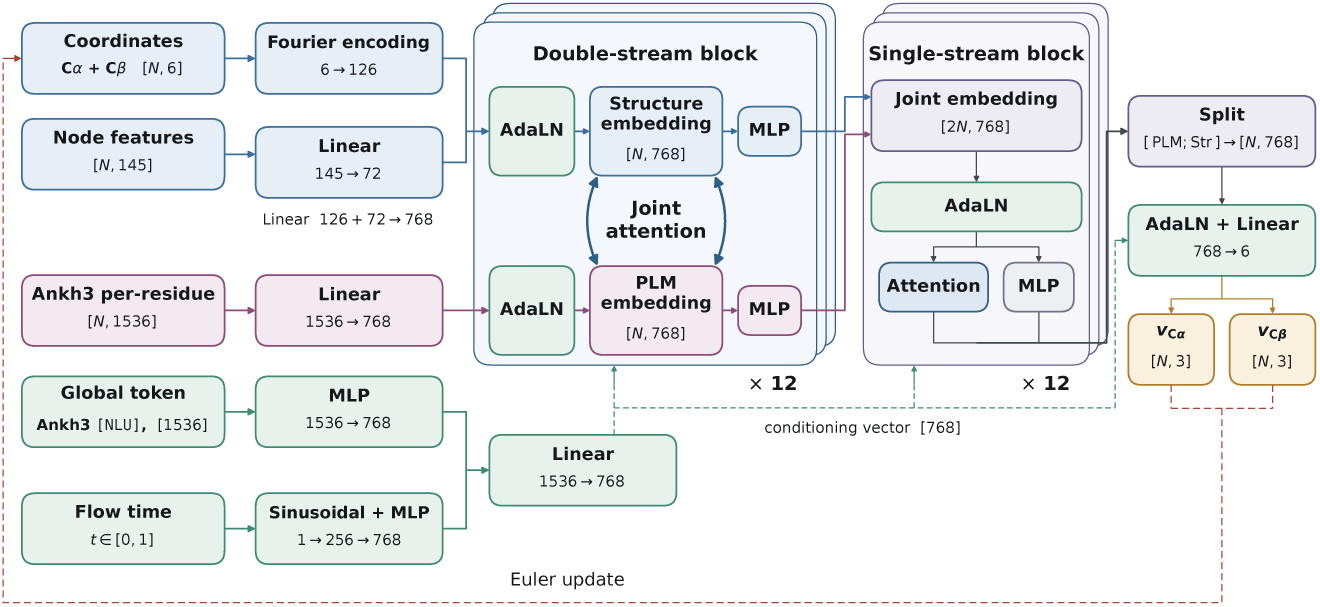
The network for a protein of *N* residues. Interpolated C*α* and C*β* coordinates are Fourier-encoded and concatenated with projected node features to form the structure stream, and frozen Ankh3-large per-residue embeddings form the sequence stream. The flow time and the global token are combined into one conditioning vector, which sets each block’s shift, scale and gate and the output head’s shift and scale. 12 double-stream blocks, which keep independent weights per stream and couple the two only inside joint attention, are followed by 12 single-stream blocks over the concatenated 2*N* tokens. Without ligand input, the network is trained toward the one holo deposition of each training pair, and 10 Euler steps of its velocity give one predicted structure. The complete diagram is Figure A2.

### Training objective and benchmark

The objective combines four structure terms (Appendix A.4),

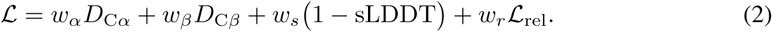

*D*_C*α*_ and *D*_C*β*_ are mean per-residue distances after one global Kabsch superposition, optimal in the least-squares sense (Kabsch, 1976; 1978). sLDDT is a smooth lDDT (Mariani et al., 2013) on C*α*, and L_rel_ scores how each residue moves relative to its neighbours in space and in sequence. The weights are *w_r_* = 2.0 and *w_α_* = *w_β_* = *w_s_*= 0.5. A2Holo was trained to minimise Equation 2 with AdamW (Loshchilov & Hutter, 2019) on the 1291 training pairs of the apo–holo collection of Peng et al. (2022), filtered and split by Somnath et al. (2023) (Appendix A.5). Training stops early on the mean validation C*α* RMSD, after 30 validations without improvement or at epoch 700 (Appendix A.6). On the 150 test pairs the released model reaches a median C*α* RMSD of 0.809 Å, against 4.46 Å for the untouched apo input and 2.12 Å for the strongest published baseline, which draws one sample per trained model (Table 1). TM-score is absent from that literature and is computed here. On this score, A2Holo beats the untouched apo input on 120 of the 150 pairs (Appendix A.8). Against each stochastic baseline’s best of 10 target-selected samples, A2Holo’s single trajectory scores higher on at least 144 of 150 pairs (two-sided Wilcoxon signed-rank, *p* ≤ 6.7 × 10*^−^*^25^, Figure A3), and on 137–144 across the 5 seeds (one pair in Figure 3). FDBM’s best of 10 costs 15.78 s per pair on one GPU, against 0.456 s for A2Holo’s single trajectory including its sequence embedding (Appendix A.10). Appendix A.9 repeats the comparison on a protein-disjoint re-split with every method retrained, where A2Holo has a lower C*α* RMSD than each baseline in 38–43 of the 58 test pairs.

**Figure 3:**
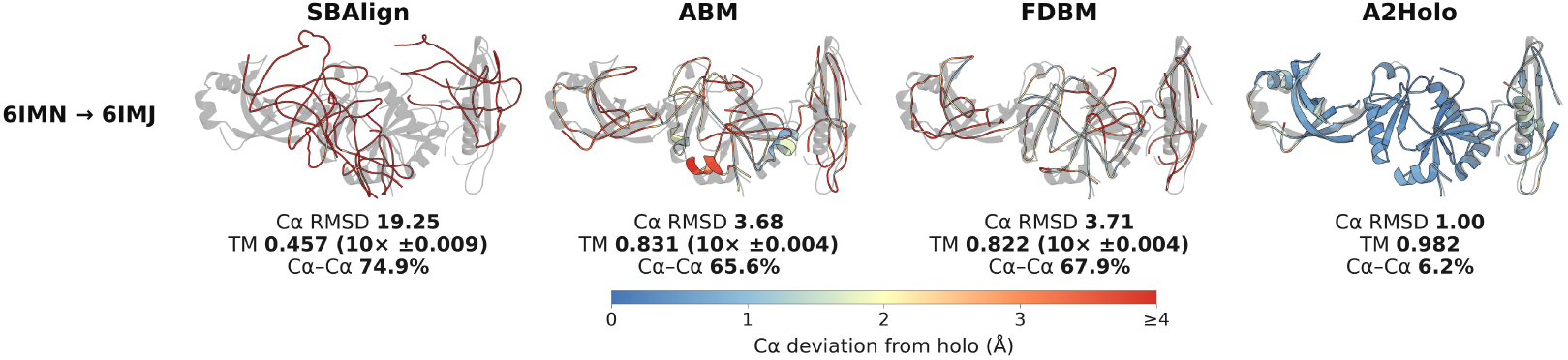
One of the hard test pairs (holo in grey). The baselines keep the best of 10 target-selected samples. A2Holo runs one trajectory. C*α*–C*α* is the share of consecutive C*α*–C*α* distances that are off the peptide length. Figure A4 shows more pairs.

Removing the sequence stream and retraining from scratch under this recipe raises the median C*α* RMSD by 0.195 Å (Appendix A.14), more than the 0.075 Å seed spread (Table A4). The ablated model also drops the double-stream blocks. Therefore, the 0.195 Å includes the loss of both. Appendix A.15 separates sequence information, fusion and model size on the protein-disjoint split.

## 4 Experiments

### 4.1 Setup and notation

Measurements are on the 9 independently trained models described below, which span 5 depths, over 133 test pairs (the 150 after a chain-length cap). They are taken at *t* = 0.5, where the structure stream is given (1 − *t*) apo + *t* holo, the interpolant the model is trained on. Intervals are per-pair bootstraps (Appendix A.11). At inference A2Holo is given the apo structure alone. Repeated at *t* = 0, 14 of 15 sign tests in this section return the same *p*, and no ordering over the 9 trainings shifts by more than 1.

Let *A*^(*ℓ*)^ ∈ R^2*N*^*^×^*^2*N*^ be the joint attention map of double-stream block *ℓ* for a protein of *N* residues, averaged over attention heads, with blocks indexed from *ℓ* = 0. Tokens are ordered [sequence; structure], so *A*^(*ℓ*)^[:*N,* :*N*] is the sequence-to-sequence quadrant and *A*^(*ℓ*)^[*N* :*, N* :] the structure-to-structure quadrant (Figure 4, five pairs in Figure A5). Each row is normalised over all 2*N* keys, and *A*^(*ℓ*)^[*q, k*] is how much query *q* reads from key *k*. For a pair, Δ*D_ij_*is the change from apo to holo in the C*α* distance between residues *i* and *j*. The *moving residues* are the top 15% by total distance change *_j_* |Δ*D_ij_*|, not by displacement |Δ**x***_i_*|. Overlap with them is the intersection over union of two top-15% sets, quoted as a multiple of the chance level, the overlap a randomly chosen 15% would reach, *f/*(2 − *f*) = 0.0811 at *f* = 0.15. Figure 4 shows these maps on one pair. In the first block the structure queries read almost only the sequence stream. In blocks 3 and 7 the structure-to-structure quadrant builds bright columns, residues that many structure queries read, and they sit at the residues that move. Section 4.2 measures this column view, the attention a residue receives, over all pairs and trainings. Figure 5 draws, along the path from apo to holo, what a structure residue receives and emits, what the sequence stream pays to it, and what the sequence stream pays within itself. On its pair, received attention is bright on the residues that move, already at *t* = 0, while emitted attention stays close to uniform, and Section 4.2 tests this contrast over all trainings. What the sequence stream pays to the structure residues is bright on one of the two moving regions, while what it pays within itself is scattered, and Section 4.4 tests that contrast.

**Figure 4:**
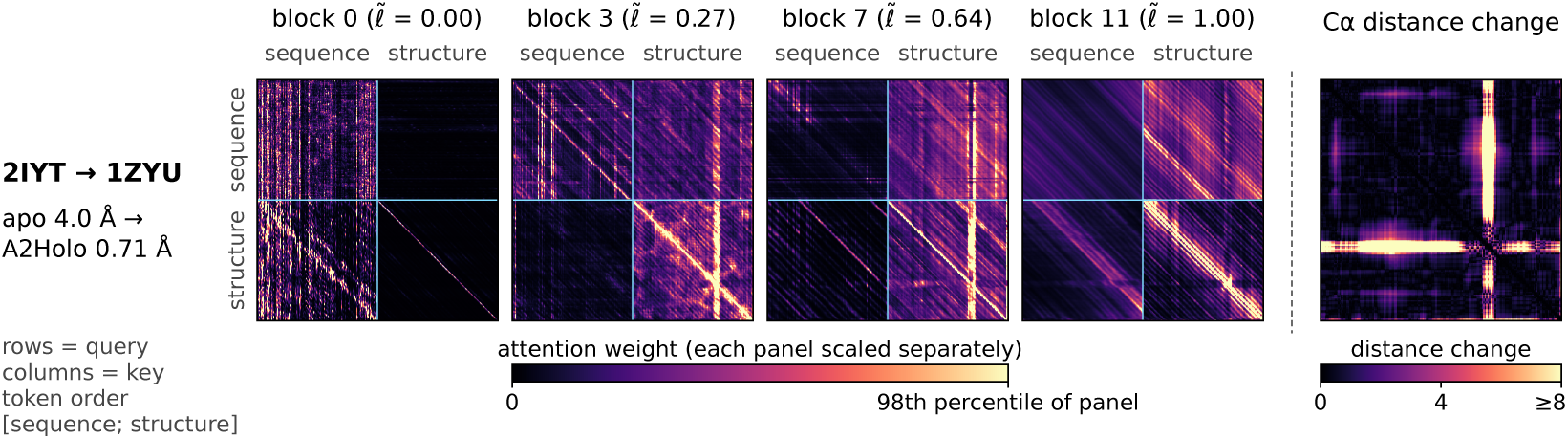
The joint attention map that Section 4 measures, for one test pair and the released model at *t* = 0.5, at four double-stream blocks. The last panel is the pair’s C*α* distance change. The pair is chosen for legibility, Figure A5 shows five pairs, and claims are made over the 9 trainings and 133 test pairs (Figure 6).

**Figure 5:**
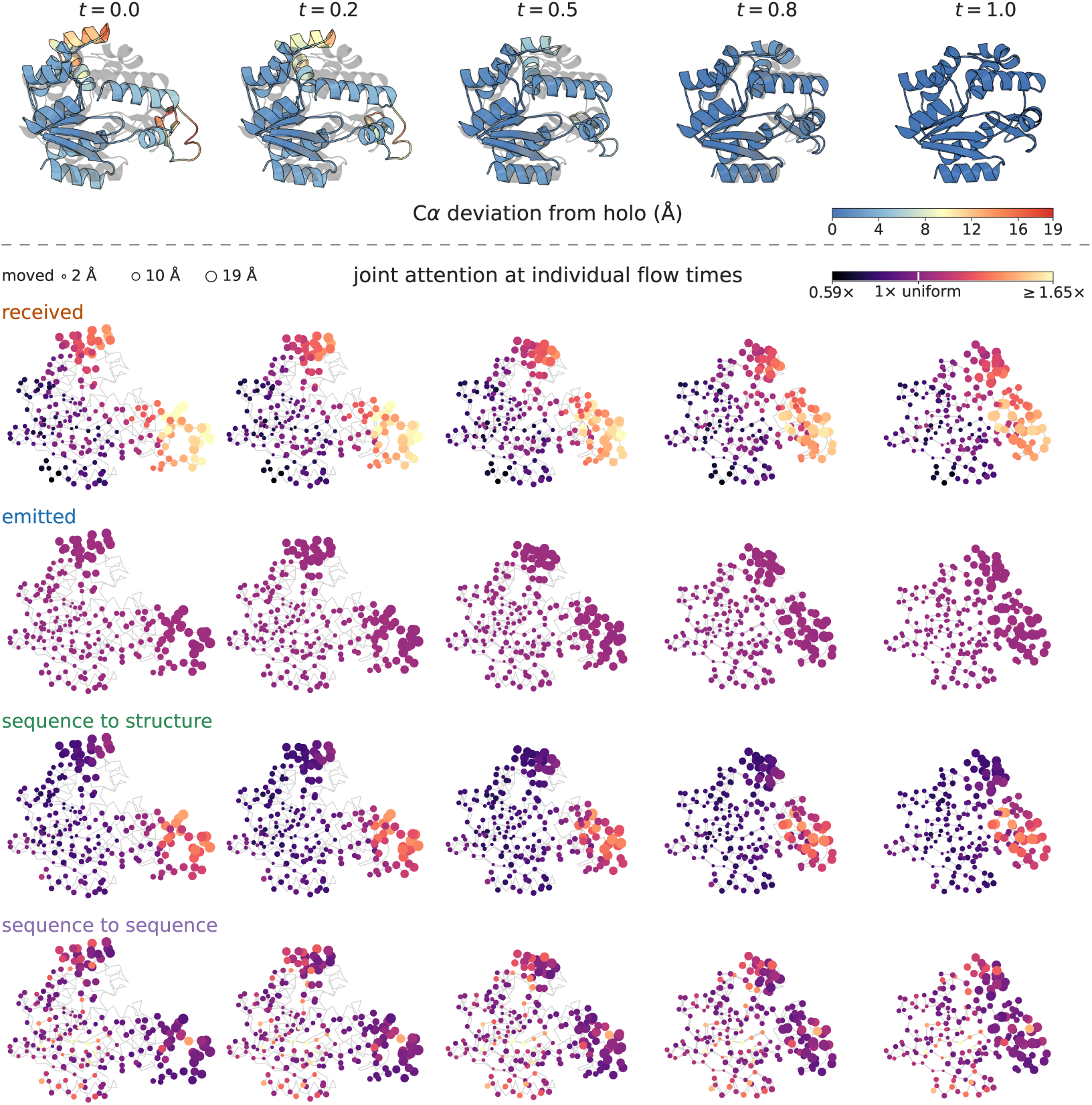
Attention on the structure, along the path from apo to holo. One test pair and the released model, at five flow times. **Top**, the structure stream’s input (1 − *t*) apo + *t* holo, each residue coloured by how far its C*α* still sits from the holo deposition, drawn over that deposition in grey. **Below**, the same five states as points, one per residue, joined by their own chain and drawn over the holo trace, both in grey. Point area is how far the residue travels. Each row colours the residues by one reduction of the joint map, block-averaged, relative to uniform attention and on one scale, clipped at its 99th percentile. In order, the rows show what a structure residue *receives* within the structure stream, what it *emits* there, what the sequence stream pays to it and what the sequence stream pays within itself. Of the 15% of residues whose distances change most, the top 15% by received attention recovers 29 of 32 at *t* = 0.5 and the top 15% by emitted recovers 12, where two unrelated sets of that size would share about 5. At *t* = 1 the input is the holo structure, so the last column marks the end of the path. The pair is chosen for legibility, and Figure 6 gives received against emitted attention over all 9 trainings and 133 test pairs.

Every claim is evaluated across the trained models rather than at a selected layer of one checkpoint. They are the released checkpoint, four further seeds of its configuration, and four trained at neighbouring depths, so a layer index means different things in different members. Nor does layer position reproduce, since the peak of the received overlap sits anywhere from 0.333 to 0.909 of the double-stream stack across the 9 trainings. On the released checkpoint, choosing the best layer per pair raises the mean overlap from 4.19× to 5.57× chance, a gain the selection produces and the model does not. Every quantity below is therefore one of five kinds: a depth average, a ratio against the chance level *f/*(2 − *f*) above (Appendix A.12), a sign-tested ordering (Appendix A.11), a count over all model × layer cells, or a curve against normalised depth *ℓ̃* = *ℓ/*(*L* − 1) over the *L* blocks of the stack being measured. A maximum or minimum over depth is quoted only as a magnitude, except at the first double-stream block, which every member shares, and as the spread of a location across trainings.

### 4.2 The emitted and received split is learned

Fix one quadrant *Q* ∈ R*^N×N^* of *A*^(*ℓ*)^, its rows indexed by the residue of the query token and its columns by the residue of the key token, and write

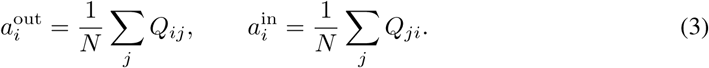

We call 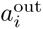 the *emitted* attention of residue *i*, which is what it reads from the key stream, and 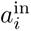 the *received* attention, which is how much the query stream reads from it. In Figure 4, a residue’s received attention is the brightness of its column in a quadrant, and its emitted attention that of its row. Per-residue attention is often summarised by adding the row and the column, which mixes the two directions (Appendix A.12).

Rows are normalised over all 2*N* keys, and the first *N* tokens are the sequence stream. Therefore, what a residue emits within its own stream and what it emits across to the other add to a fixed total. For a structure-stream query,

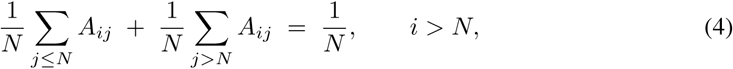

with the two roles swapped for a sequence-stream query. The two emitted vectors of a stream sum to 1*/N* at every residue, so *ρ*_cross_ = − *ρ*_intra_ against any fixed target and reporting both counts one measurement twice. Column sums carry no such constraint.

Separating the two quantities of Equation 3 shows that their sum mostly measures the received direction. Depth-averaged over the 9 models, structure-to-structure attention overlaps the moving residues at 1.88–2.54× chance when measured on what a residue receives. Measured on what it emits, the overlap is 1.03–1.75× (Figure 5, first two rows, on one pair), and summed it is 1.78–2.50×. Training by training, the summed value sits on average 0.07× chance from the received one and 0.71× from the emitted one. Within every training the received value exceeds the emitted one (9 of 9, sign test *p* = 0.0039, Figure 6). At cell level, received exceeds emitted with disjoint bootstrap intervals in 56 of the 108 model × layer cells, against 19 the other way. Emitted attention is much weaker than received but not absent, clearing chance in 51 of 108 model × layer cells against 85 for received.

**Figure 6:**
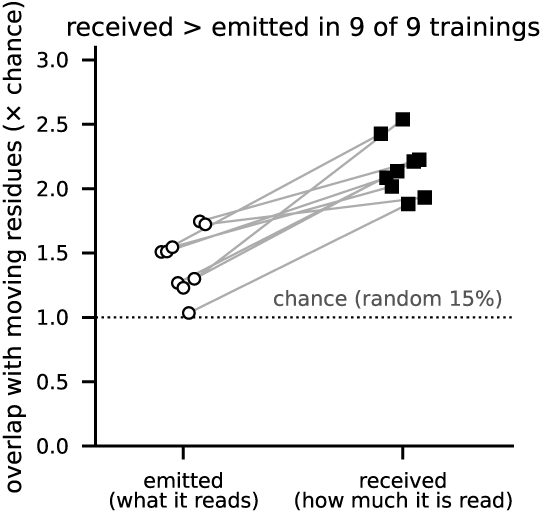
Received attention picks out the moving residues better than emitted attention, in every training. One line per training, depth means of the overlap over 133 pairs at *t* = 0.5 (Figure A6 by depth).

At random initialisation, with architecture, inputs and measurement unchanged, the received-to-emitted ratio averages 0.94–1.02 across 8 depths, against 1.12–2.06 trained, and every training beats the mean of the 3 initialisations at its own depth (9 of 9, *p* = 0.0039), so the asymmetry is learned and is not row-normalisation bookkeeping. A second control changes neither the architecture nor the seed, but what the model was trained to do. With the flow time drawn uniformly in place of the recipe’s distribution, the same configuration returns a ratio of 0.970, inside the untrained band. Placing the mass near the holo end instead gives 1.322, which keeps the asymmetry. A trained model can therefore lack the split altogether, so it depends on the objective and does not follow from the layout alone.

### 4.3 Attention picks out moving residues beyond an apo-centroid baseline

An apo-only geometric rule, distance from the apo centroid, recovers the same moving residues at 3.72× chance. That is above the 1.88–2.54× of depth-averaged received attention, so absolute overlap cannot separate attention from that rule. Holding centroid distance fixed does. Partialled on it, the rank correlation between received attention and total distance change runs 0.046 to 0.145. Untrained networks of the same depth give depth means of 0.004 to 0.012, and every training exceeds the mean at its own depth (9 of 9 trainings, *p* = 0.0039). For emitted attention the correlation runs −0.144 to −0.005, and received exceeds it (9 of 9, *p* = 0.0039). Restricting instead to the residues the rule leaves unselected gives the same ordering, received 1.51–2.20× chance against emitted 1.03–1.55× in 9 of 9 (*p* = 0.0039). The remaining residues still carry the rule’s ranking, so only this ordering, not its size, is claimed.

Taken over residue pairs (*i, j*) rather than single residues, the structure-to-structure map barely correlates with |Δ*D*| (Pearson *r*, entry by entry). The depth-mean correlation is positive in 8 of 9 trainings (*p* = 0.039) but reaches only −0.046 to 0.042, and in 9 of 9 trainings the block-level values carry both signs. Attention marks where the protein changes, not how it changes.

### 4.4 What crosses between the streams

The share of attention mass that crosses is 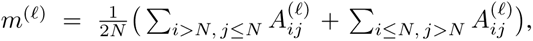 for which uniform attention gives *m* = 1*/*2. Depth-averaged per model, *m* sits at 0.434–0.537 across the 9 trainings, so the two streams divide their attention almost evenly. The signal sits at the structure-residue end of the cross quadrants, resolved in the sequence-to-structure one. The attention a structure-stream residue pays to the sequence stream overlaps the moving residues at 1.35–3.10× chance (Table 2). By Equation 4 it is fixed by the emission Section 4.2 reports, so it adds no information beyond that vector. The attention the sequence stream pays to a structure-stream residue overlaps them at 1.93–2.82×, and that one is independent. It exceeds the sequence stream’s own internal received attention (1.12–1.86×) in 9 of 9 trainings (*p* = 0.0039), as the last two rows of Figure 5 show on one pair. No quantity indexed by a sequence token exceeds 1.87× chance, and received attention indexed by a structure residue stays above all four of them in every training (9 of 9, *p* = 0.0039, Table 2). At random initialisation the same quantities sit at 0.96–1.26× with no such separation, and the ordering holds in only 2 of 24. Quadrant by quadrant the structure end beats the sequence end in 9 of 9 trainings in the sequence-to-structure quadrant (*p* = 0.0039) and in 7 of 9 in the structure-to-sequence one, which this test cannot resolve.

**Table 2:** Overlap with the moving residues in each quadrant of the joint attention map, as a multiple of the chance level *f/*(2 − *f*), depth-averaged within a training and ranged over the 9 trainings at *t* = 0.5. Brackets give 24 random initialisations. A dagger marks the entries a structure residue indexes, that is, the column of a received quantity and the row of an emitted one.

| query → key | emitted (row) | received (column) |
|---|---|---|
| structure → structure | **1.03–1.75**<sup>†</sup> (1.04–1.24) | **1.88–2.54**<sup>†</sup> (1.03–1.22) |
| sequence → structure | 1.05–1.63 (1.03–1.17) | **1.93–2.82**<sup>†</sup> (1.03–1.22) |
| structure → sequence | **1.35–3.10**<sup>†</sup> (1.02–1.32) | 1.29–1.87 (1.00–1.26) |
| sequence → sequence | 1.00–1.46 (0.96–1.18) | 1.12–1.86 (1.03–1.24) |

At moving residues the structure stream both reads more from the sequence stream and is read more by it. Neither exceeds the other consistently (4 of 9, *p* = 1.00), so neither cross-attention direction dominates.

The size of each residue’s displacement is tracked as well. Averaged over depth, the rank correlation between how much a structure-stream residue is read by its own stream and |Δ**x***_i_*| runs −0.056 to +0.143, positive in 8 of 9 trainings (*p* = 0.039). What that residue reads from the sequence stream runs −0.060 to +0.215 and is positive in 6 of 9 trainings (*p* = 0.51), which this test cannot resolve. The emitted intra-stream correlation mirrors that cross-stream reading by Equation 4 and is not shown.

### 4.5 Computation across the stack

The layout splits depth into a double-stream and a single-stream stack, a split inherited from the image setting. With one block’s residual update removed at a time, a double-stream block costs more than a single-stream one in 8 of 9 trainings (Figure A7, Appendix A.16). Because every double-stream block precedes every single-stream one, block type and depth are confounded in this comparison.

Damping only the cross-stream quadrants separates joint attention from the block that holds it. Zeroing the two cross-stream quadrants of one double-stream block at a time raises the one-step C*α* RMSD by 0.322–1.036 Å (depth mean, Appendix A.16), more than zeroing the two intra-stream quadrants, 0.244–0.732 Å, in 8 of 9 trainings (*p* = 0.039). That intervention does show the response is non-linear. Halving a quadrant’s weight costs 0.004–0.019 Å against 0.322–1.036 Å for zeroing it. Rows are renormalised after the intervention, so zeroing a quadrant routes its mass to the other rather than deleting it.

Block removal asks which blocks the prediction depends on. Probing asks what each block’s representation carries. Each pair sits in its own arbitrary frame, so the probe targets are rotation-invariant (Appendix A.17). To separate what a block carries from a flexibility prior already fixed by sequence, we measure decodability *relative to a sequence-only baseline*,

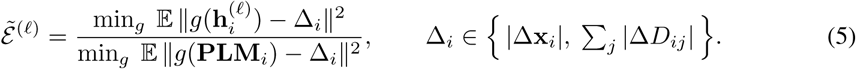

Here *g* is an affine map fitted by ridge regression on residues of the other folds, one per input, 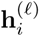 is the state of residue *i*’s structure token leaving block *ℓ*, **PLM***_i_* is the frozen language-model embedding of the same residue, and E runs over held-out residues (Appendix A.17). A value below one means the target is more decodable from block *ℓ*’s state than from the language-model embedding alone.

Averaged over depth, Ẽ runs 0.320–0.569 for displacement magnitude and 0.549–0.845 for total distance change. It is below one for both targets (9 of 9 trainings, *p* = 0.0039, Figure A8). Where the per-model minimum over depth sits does not reproduce. The block with the lowest Ẽ falls anywhere from *ℓ̃* = 0.348 to 0.957 over both stacks for displacement, and from 0.348 to 0.935 for total distance change. At the first double-stream block, the one position every member shares, Ẽ is already below the baseline in 9 and 8 of 9 trainings, at 0.526–0.985 and 0.736–1.035. It falls further after the first block, so the advantage is not handed over by the input alone. Nor does it come from the architecture. Untrained networks of the same architectures give depth means of 1.071–1.204 and 1.034–1.187. Ẽ lies below the mean of untrained networks of the same depth for both targets (9 of 9 trainings each, *p* = 0.0039, Appendix A.17).

## 5 Conclusion

A2Holo takes the dual-stream joint attention of in-context image generation and applies it to apo-to-holo transport. On the D3PM benchmark one deterministic trajectory, given no ligand and drawing no samples, reaches a higher TM-score than every stochastic baseline’s best of 10 target-selected samples on at least 144 of 150 pairs (Figure A3).

The transplanted mechanism was then measured rather than assumed. Across independently trained models of 5 depths, the answer is that the structure stream is read where the protein moves. The attention a residue receives marks the moving residues more than the attention it pays, and this split is learned. The received signal survives holding distance from the apo centroid fixed (Section 4.3). The sequence stream’s attention to the structure stream also picks out those residues better than its attention to itself (in every model). Along the stack the structure stream carries more about the coming change than the language model alone, and all of this reproduces across models and with the apo structure alone as input, while the depth at which it peaks does not. Attention marks where the protein changes, not how. On the protein-disjoint split, pairs whose structure-stream attention marks the moving residues more closely show less off-target motion (Spearman −0.42 over 58 pairs, Holm-adjusted *p* = 0.035, Appendix A.9). A2Holo therefore provides not only a deterministic route from an apo structure to a holo one, but a measured account of how the two modalities are combined along that route. That account is possible because both streams run over the same residues, so every entry of the joint map belongs to a residue pair, which text and image tokens in the original setting do not share.

All results come from one benchmark, and those of Section 4 from 9 trainings of this layout, so they describe what this layout learned there rather than a general property of proteins or of joint attention.

The model itself outputs only C*α* and C*β* positions, commits to one deterministic state and sees no ligand (Appendix A.19). Each of these is a direction for future work. Full-atom models could come from torsional side chains or from giving the predicted C*α* trace to a structure predictor as a template (Passaro et al., 2025). Multi-state output would cover proteins with more than one bound form, and ligand conditioning would tell the model what the pocket will hold. Docking-protocol evaluation (Morehead et al., 2026b) would then test the result in use, and apo-to-holo transition generation may become practical for structure-based design.

## AI USE STATEMENT

In this work, we used generative AI tools, specifically large language models, for the code, for the figures and for editing the prose. They assisted in writing the code used to train the model and to analyse its outputs, and in producing the figures from those outputs. The text of this manuscript was written by the authors, and AI tools helped revise and polish it. We did not use them to generate synthetic datasets or to prove any mathematical claim.

We have reviewed all AI-assisted work. The authors read and tested the AI-assisted code before any of its output entered the manuscript. They checked every passage an AI tool revised against the results it reports. We take responsibility for the final content of this work, including text, claims or artifacts produced with the aid of generative AI.

## Ethics statement

This work predicts the conformational change of a protein’s C*α* and C*β* positions from experimentally determined structures that are already public. It involves no human subjects and no personal or otherwise sensitive data, and it releases no new dataset. All structures are Protein Data Bank entries, from the D3PM database (Peng et al., 2022) as filtered by Somnath et al. (2023). The model moves the C*α* and C*β* positions of a protein that already exists. It designs no molecule, no sequence and no binder, so it adds no design capability of its own, and its intended use is as a component of structure-based drug design pipelines. The limitations of the method in that role are stated in Section 5 and Appendix A.19, and we regard overstating its reliability as the principal foreseeable harm.

## Reproducibility statement

Full methods and supplementary analyses are given in Appendix A. Within it, the problem formulation is in Appendix A.1 and the dual-stream architecture in Appendix A.3. The training protocol, with the deployed model’s hyperparameters, its optimiser schedule and its stopping criterion, is in Appendix A.6. The deterministic inference procedure is in Appendix A.7 and the statistical tests are in Appendix A.11. Inference injects no noise and does not sample time, so a given checkpoint returns the same structure on every run and reproduces a reported value up to floating-point precision rather than in distribution. Retraining does not. The same configuration trained 5 times with different seeds spans 0.761–0.836 Å in median C*α* RMSD (Appendix A.13). We use the 1591 apo–holo pairs of the D3PM database (Peng et al., 2022) as filtered by Somnath et al. (2023), with their 1291/150/150 training, validation and test split taken unchanged so that published values remain comparable (preprocessing in Appendix A.5). The evaluation protocol is in Appendix A.8, including which baseline numbers are literature-reported rather than reproduced by us and which were obtained by running the checkpoints those baselines released.

The attention analysis is made over 9 independently trained models rather than one checkpoint, so a reported effect is not a property of a single training. The 9 are 5 trainings of this configuration differing only in the random seed and 4 of different depth trained at seed 1.

The training and analysis code is available at https://github.com/ding-oh/A2Holo. The repository also links the trained checkpoints. There are eight, the released model and the seven of the protein-disjoint comparison in Appendix A.15, and each carries a SHA-256 so that a reader can confirm which weights they hold.

## Author Contributions

D.K. and J.L. conceived the study. D.K. and H.K. developed the methodology. D.K., J.S., M.K. and M.J. carried out the experiments, and H.K. performed the formal analysis. D.K., H.K., J.S., M.K. and M.J. validated the results. D.K. and H.K. prepared the figures and wrote the original draft. D.K., H.K. and J.L. reviewed and edited the manuscript. J.L. supervised and administered the project, provided resources and acquired funding.

## Acknowledgments

This work was supported by the Institute of Information & Communications Technology Planning & Evaluation (IITP) grant funded by the Korea government (MSIT) (RS-2023-00220628, Artificial intelligence for prediction of structure-based protein interaction reflecting physicochemical principles). It was also supported by two National Research Foundation of Korea (NRF) grants funded by the Korea government (MSIT), RS-2024-00352229 and RS-2026-25506638 from the Bio&Medical Technology Development Program, and by the NRF Basic Science Research Program funded by the Ministry of Education (RS-2026-25563184). Further support came from the Korea Machine Learning Ledger Orchestration for Drug Discovery Project (K-MELLODDY, RS-2026-25608347), funded by the Ministry of Health & Welfare and the Ministry of Science and ICT, and from the Korean ARPA-H Project through the Korea Health Industry Development Institute (KHIDI, RS-2026-25614921), funded by the Ministry of Health & Welfare, Republic of Korea. Additional support came from the Samsung Science & Technology Foundation (C1016338-01-01) and from the Korea Ministry of Environment through the Chemical Accident Prevention Technology Development Project (RS-2023-00219144). Finally, Seoul National University supported this work through the AI-Bio Research Grant (0413-20230053) and provided computing resources on its cluster.

## A Supplementary methods and analyses

### A.1 Problem formulation

Section 3 poses the task as noise-free transport along the straight path **x***_t_* = (1 − *t*) **x**_apo_ + *t* **x**_holo_, from the apo state at *t* = 0 to the holo state at *t* = 1. Supervision is applied to the position the velocity implies, **x**^_holo_ = **x**_apo_ + **v***_θ_*(**x***_t_, t*), rather than to the velocity itself. C*α* and C*β* are interpolated independently and predicted in the same forward pass, so the model solves two three-dimensional flows that share a frame only through the loss.

### A.2 Inputs and their projections

Five inputs enter the network, and each is projected separately. The current C*α* and C*β* coordinates give 6 channels per residue, which a Fourier feature encoding with 10 frequencies expands to 126 dimensions with the raw input retained (Tancik et al., 2020). In parallel, a learned linear map compresses 145 scalar node features to 72 dimensions. Those features are an amino-acid one-hot code and physicochemical descriptors inherited from the benchmark preprocessing. The two are concatenated to 198 dimensions and projected to the model width, forming the structure stream. Residue identity thus enters both streams. Only the structure stream carries the current conformation.

Per-residue sequence representations are taken from the Ankh3-large encoder and give 1536 dimensions per residue. They are extracted once, from the sequence recovered from the node features, and frozen thereafter (Alsamkary et al., 2025). Recovering the sequence that way guarantees a one- to-one correspondence between embedding and residue. The global token is the encoder output at position 0, which is the `[NLU]` mode prefix that Ankh3 prepends to the sequence. Ankh3 itself mean-pools for sequence-level tasks, and A2Holo instead reads the position-0 output as its whole-sequence summary. Figure A1 measures how much the embedding and the apo structure already carry of each other.

Projections are deliberately non-uniform. Each per-residue stream enters through a single linear layer, whereas the two global inputs, the flow time and the global token, enter through two-layer MLPs, following the image backbone the architecture is adapted from (Black Forest Labs et al., 2025; Esser et al., 2024).

### A.3 Dual-stream architecture and conditioning

The network stacks 12 double-stream and 12 single-stream blocks of width 768, with 8 attention heads and an MLP ratio of 4.0 (Figure A2). Section 3 describes how the blocks couple the two streams, and the details it leaves out are these. In a double-stream block the sequence stream comes first along the token axis, so the structure half is the second half of the output. Queries and keys are normalised before attention (Dehghani et al., 2023) with RMSNorm (Zhang & Sennrich, 2019), and the one-dimensional rotary embeddings use a base of 10,000. In a single-stream block the parallel attention and feed-forward branches are rejoined by one linear layer. A final adaptive layer normalisation and linear map return 6 channels, split into the C*α* and C*β* velocities. Both global inputs reach the network through a single conditioning vector, formed by concatenating the time and global token embeddings, layer-normalising them and projecting to the model width. Each block derives its own shift, scale and gate parameters from this vector, and these modulation parameters are not shared between blocks (Peebles & Xie, 2023).

**Figure A1:**
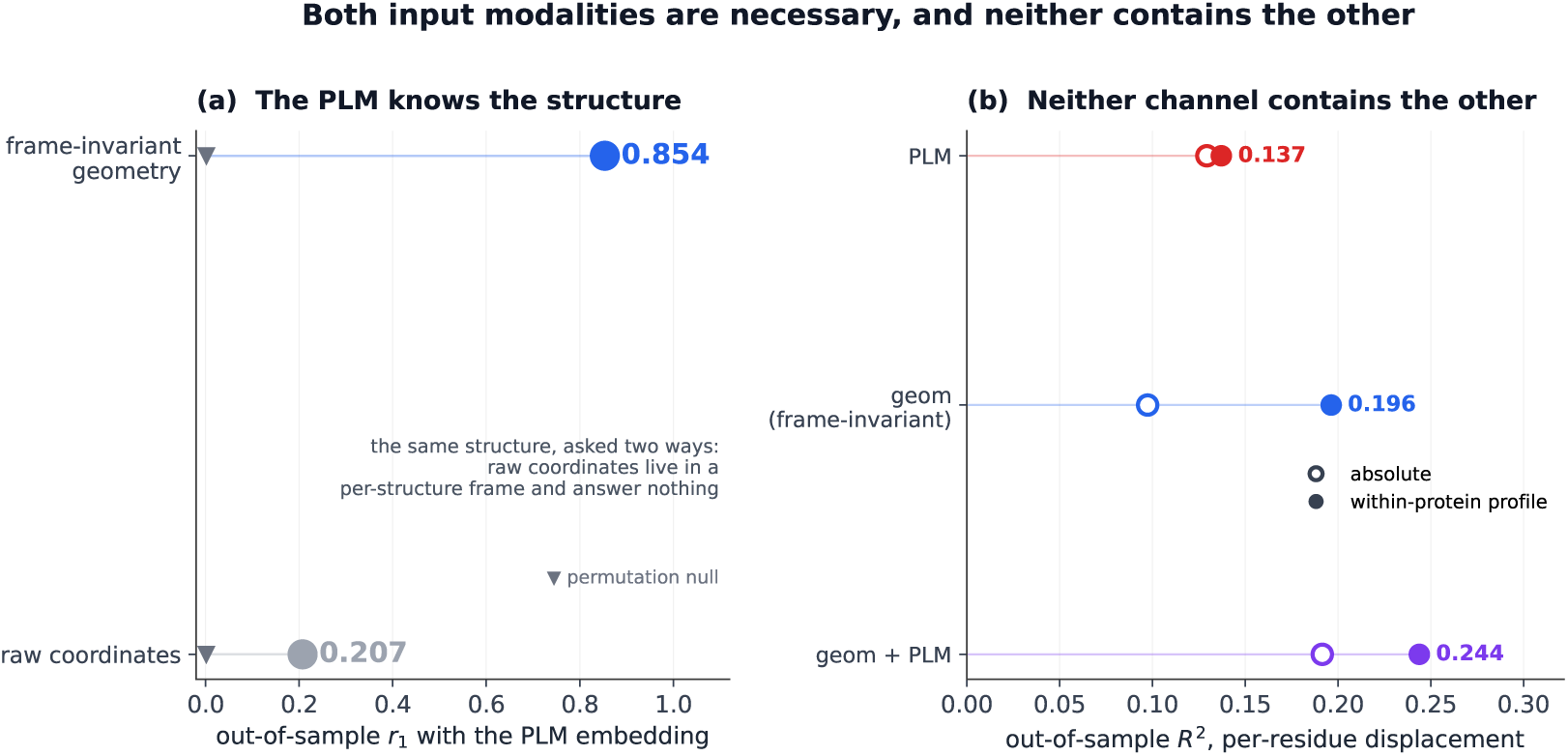
The language-model embedding and the apo geometry do not contain each other, measured on the inputs alone with no trained model. Statistics are out-of-sample over 2100 apo–holo pairs (698231 residues), every processed pair of the D3PM collection with an embedding, before the filter that leaves the 1591-pair benchmark. **a**, Top canonical correlation between the Ankh3 embedding and the apo structure. Against rotation- and translation-invariant descriptors (contact numbers, burial, pseudo-torsions, local radius of gyration) it reaches *r*_1_ = 0.854, so a sequence-only representation already carries much of the fold, as far as a linear map can see. Against the physicochemical features often concatenated beside the embedding, the leading canonical correlation is 0.995. Against raw C*α* and C*β* coordinates it falls to 0.207 (descriptors against coordinates: 0.063), an effect of each structure’s arbitrary frame. Triangles are permutation nulls. **b**, Out-of-sample *R*^2^ for predicting per-residue displacement from each channel. Open markers, absolute displacement. Filled markers, the within-protein profile. On the profile, apo geometry leads (0.196 against 0.137), and together the two reach 0.244, above either alone. These are linear probes, so they bound only linearly accessible information, which suffices for the non-redundancy they show.

**Figure A2:**
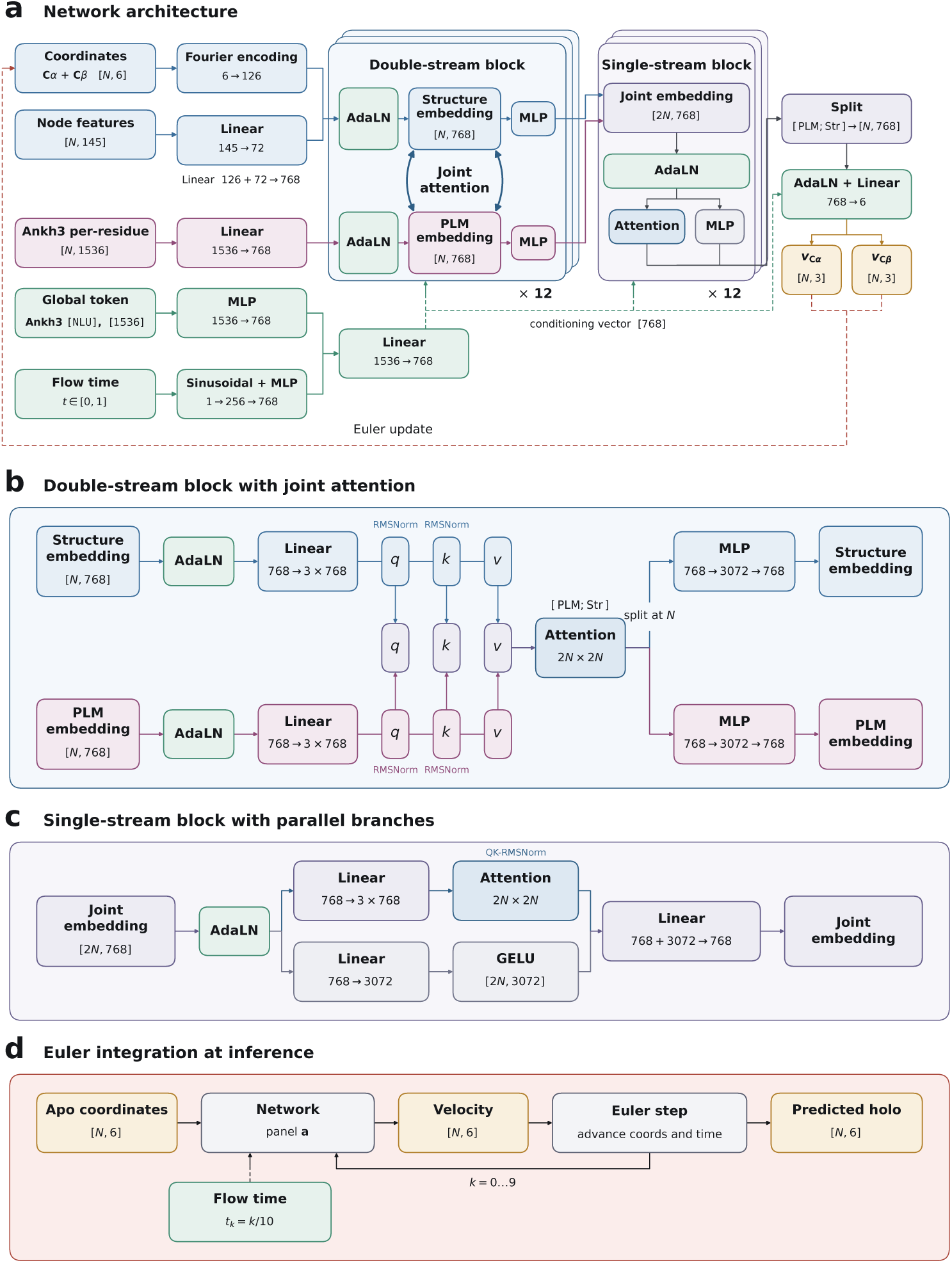
The complete architecture diagram. Figure 2 shows panel **a** alone. **a**, The whole network for a protein of *N* residues. The structure stream is 768-dimensional, and its inputs and their projections are described in Appendix A.2. **b**, One double-stream block with joint attention. Queries, keys and values are formed per stream and concatenated along the token axis *with the sequence stream first*, attended over once as a single 2*N* × 2*N* map, then split back at *N*. The two streams keep independent weights everywhere else. The panel labels the sequence stream PLM embedding and the structure stream Structure embedding, and abbreviates their concatenation [PLM; Str]. Every quadrant read in Section 4 is defined against that token order. AdaLN is layer normalisation whose scale and shift are set by the flow time and the global token. **c**, A single-stream block. Attention and the feed-forward branch are computed in parallel from the same modulated input and rejoined by one linear map, rather than applied one after the other. **d**, The deterministic inference loop, 10 equal forward-Euler steps from the apo coordinates, with no sampling and no injected noise at any step.

### A.4 Training objective

The objective is Equation 2, with *w_α_* = 0.5, *w_β_* = 0.5, *w_s_* = 0.5 and *w_r_* = 2.0. Each term is averaged within a protein and then over the proteins of a batch.

The two structure terms are mean per-residue distances after Kabsch superposition. Let **x̂***_i_* and **x***_i_* be the predicted and target C*α* coordinates of residue *i*, with centroids **x̄** and **x̄**. Let *R* be the rotation that best superposes the centred prediction on the centred target (Kabsch, 1976). Then

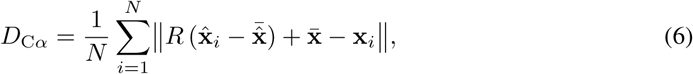

where ‖·‖ is the unsquared Euclidean norm on R^3^, here and in Equation 8, with no clamp on the per-residue distance. *D*_C*β*_ is Equation 6 with the same *R*, **x̄**^ and **x̄** applied to the C*β* coordinates and averages over the residues with a real C*β*. Glycine is therefore masked, and the two channels are scored in one frame.

The third term is a smooth, differentiable lDDT (Mariani et al., 2013) on C*α*. Let *d_ij_*and *d*^^^*ij* be the target and predicted distances between residues *i* and *j*, and P = {(*i, j*) : *i* /= *j, d_ij_ <* 15 Å}. Then

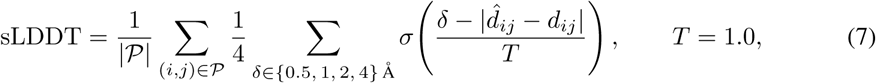

with *σ* the logistic sigmoid. It needs no superposition.

The fourth term scores how each residue moves relative to its neighbours, which one global super-position barely constrains. Write **v***_i_* = **x̂***_i_* − **x**_apo*,i*_ for the predicted C*α* displacement of residue *i* and **v***^∗^* = **x***_i_* − **x**_apo*,i*_ for the true one. Each residue is joined to its 16 nearest residues in the apo structure and to the 2 residues on either side of it in sequence, which fixes a neighbour list E for each protein. Then

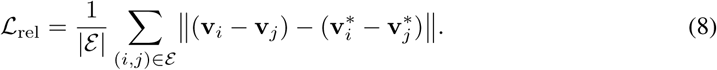

It is computed without superposition and is unchanged by a translation of the whole field. The neighbour list comes from the apo coordinates and carries no gradient. Bond-length and steric-clash terms are implemented and were held at zero weight.

### A.5 Dataset and splits

Training and evaluation use the D3PM collection of apo–holo pairs as curated for the SBAlign benchmark, comprising 1591 pairs divided into 1291 for training, 150 for validation and 150 for testing. We take the split unchanged, so that values published on it remain directly comparable with ours (Somnath et al., 2023; Peng et al., 2022). Appendix A.9 evaluates A2Holo retrained on a protein-disjoint re-split of the same pairs.

Before training, apo C*α* was Kabsch-superposed on holo C*α*, and each structure was translated to put its own C*α* centroid at the origin, so the displacement Δ**x***_i_* averages to zero over the residues of every pair. Test pairs are prepared the same way. Rotating the apo input at random moves the predicted shape by a median of 0.075 Å (Appendix A.8), so the prediction does not rest on that shared orientation. C*β* atoms were extracted from the deposited structures, with glycine receiving a virtual C*β* placed from its N, C*α* and C atoms at ideal backbone geometry. A per-residue mask indicates which C*β* positions are real. The C*β* channel received the same superposition and the same shift by the C*α* centroid.

### A.6 Training protocol

Models were trained on the objective of Appendix A.4 with AdamW (Loshchilov & Hutter, 2019) at a base learning rate of 0.0001 defined for a reference batch of 32 proteins and scaled linearly with the batch actually used. The batch was fixed at 20 proteins, giving an effective learning rate of 0.0000625. Weight decay was 0.00001, gradient norms were clipped at 5.0, and no dropout was applied anywhere in the network.

The flow time is drawn per protein from a Beta(2, 2) distribution, which concentrates supervision on the middle of the path, away from its two ends. Rotation augmentation draws an independent uniform rotation for each protein in the batch and applies it to the apo and holo coordinates of both channels about the apo centroid. Rotation consistency is therefore learned rather than imposed by equivariant operators, a choice AlphaFold 3 (Abramson et al., 2024) and SimpleFold (Wang et al., 2026c) also make. Appendix A.8 measures how far it holds.

Forward and backward passes run under bfloat16 autocast, with the Kabsch superposition, the loss and the rotary phase angles computed in single precision. An exponential moving average of the weights is maintained with decay 0.9999. Validation runs every 5 epochs and integrates the flow with a single Euler step for efficiency. Its statistic is the mean C*α* RMSD over the validation pairs. The learning rate is reduced by a factor of 0.7 after 3 validations without improvement, and the checkpoint kept is the one with its lowest value. Training stops after 30 validations (150 epochs) without improvement or at epoch 700, the defaults of the released code. Replayed on the validation records of every seed and every ablation run, this rule selects the checkpoint reported here. The released model’s checkpoint is from epoch 640, and it was trained on a single GPU.

### A.7 Deterministic inference

Inference integrates the learned field forward from the apo structure with 10 forward-Euler steps of equal size (Equation 1) on a deterministic time grid. No time is sampled and no noise is injected, so the procedure returns the same single pose on every run. There is no ensembling, no best-of-*N* selection and no averaging over trajectories. Every benchmark number in Table 1 uses the exponential moving average weights and 10-step integration. The attention measurements are single forward passes, and the block-removal and quadrant-damping costs of Section 4.5 are one-step predictions at *t* = 0.5 (Appendix A.16). Ten steps beat one on 99 to 116 of 150 pairs in each of the 5 seed trainings (two-sided Wilcoxon signed-rank, *p* ≤ 1.8 × 10*^−^*^5^). Ten steps and a hundred are separated by a median of 0.0008 Å per pair, detectable on the most favourable training (*p* = 2.2 × 10*^−^*^3^) but far below the scale this task is measured at.

### A.8 Evaluation and baseline protocol

Accuracy is reported as the RMSD between predicted and experimental holo coordinates after Kabsch superposition, over the 150 pairs of the test split. C*β* RMSD is computed under the transform fitted on C*α*, with glycine excluded, so that the C*β* channel is never given an alignment of its own. Global fold agreement is measured as the TM-score returned by US-align in monomer mode for proteins, normalised by the length of the target structure (Zhang & Skolnick, 2004; Zhang et al., 2022). The apo input is scored in exactly the same way and serves as the no-prediction reference.

Baseline values enter the paper by two routes. The C*α* RMSDs of EGNN, SBAlign, Sesame, ABM and FDBM are quoted as reported in Nobis et al. (2025), Table 1, which is the one place all five appear together. That table itself draws its rows from different sources. EGNN and SBAlign come from Somnath et al. (2023) and Sesame from Miñán et al. (2025). ABM comes from a retraining the authors of Nobis et al. (2025) performed themselves, since De Bortoli et al. (2023) report no protein experiments. The rows also differ in sampling. ABM and FDBM take one sample from each trained model and average over five trainings (Nobis et al., 2025, Appendix F.2). SBAlign draws ten samples per pair and scores all of them, so its row is a typical sample rather than a selected one, and its RMSD fractions are fractions of those samples where ours are fractions of pairs. Sesame reports no sample count. TM-scores are not reported anywhere in that literature and were computed by us from the released weights on the same test split. The quoted RMSDs and the TM-scores therefore describe different runs of the same methods.

SBAlign, ABM and FDBM ship weights trained on this split. All three were re-run through the inference code released with them, without retraining. 10 samples per pair give the TM-score comparison, which reports the best of the 10. Selecting the best of 10 samples requires the target structure to select against. It is therefore an oracle a deployed user does not have, and it places every stochastic baseline at an upper bound it could not reach in practice. A2Holo uses neither the 10-sample budget nor the oracle. Its transport is an ordinary differential equation, so one integration returns the same structure every time. On TM-score, A2Holo is higher than the best FDBM sample on 144 of the 150 pairs and lower on 6 (*p* = 6.7 × 10*^−^*^25^). The counts are 145 and 5 against ABM (*p* = 1.9 × 10*^−^*^25^), 150 and 0 against SBAlign (*p* = 2.3 × 10*^−^*^26^), and 120 and 30 against the untouched apo input (*p* = 1.9 × 10*^−^*^20^). Each *p* is a two-sided Wilcoxon signed-rank test on the per-pair differences, and Figure A3 shows every pair. The quoted RMSD column admits no such test, since it is published as medians only.

**Figure A3:**
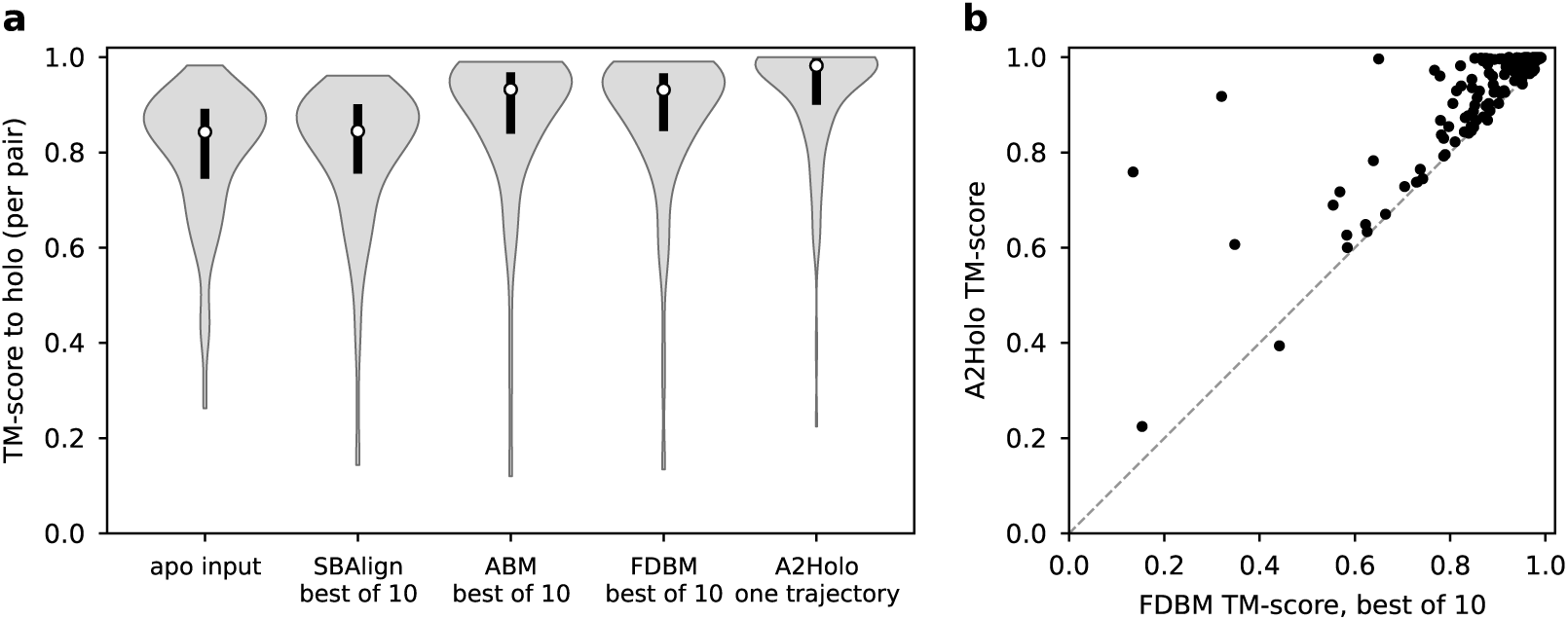
Per-pair TM-scores on the 150 test pairs. **a**, Distribution for the apo input left untouched, then SBAlign, ABM and FDBM, which keep their best of 10 samples selected using the target, and A2Holo, which integrates one trajectory. Bars span the interquartile range and open circles mark the median. **b**, A2Holo against the best FDBM sample, one point per pair. Points above the dashed diagonal are pairs where A2Holo scores higher, 144 of 150 (*p* = 6.7 × 10*^−^*^25^, two-sided Wilcoxon signed-rank).

Sesame and EGNN ship no weights usable on this split, so only their published RMSDs are available.

Figure A4 shows the structures behind these numbers on five test pairs, one per difficulty level. All four columns, and the grey holo behind them, are cg2all (CalphaBasedModel) reconstructions (Heo & Feig, 2024) from C*α* with one residue numbering per pair, because every baseline predicts C*α* only. The reconstruction is for display. All numbers in Figure A4 are computed on C*α*, the one atom every method predicts. The C*α* RMSDs printed there for the baselines are our own, computed for the TM-best of their 10 samples, and they are not the medians quoted in Table 1, which come from a different protocol. Over the 150 test pairs, the median fraction of consecutive C*α*–C*α* distances more than 0.5 Å from 3.8 Å is 0.0% for both the holo depositions and the apo inputs, 1.1% for A2Holo, and 30.5%, 33.2% and 70.2% for FDBM, ABM and SBAlign at their best of 10, with a largest single gap of 8.3 Å against 24.0, 29.8 and 41.3 Å. The unravelled appearance of the baseline columns is therefore a property of their output and not of the reconstruction, which is identical for every column. Those fractions are not normalised for the motion each method performs, which is why the untouched apo input sits beside the depositions on them. A chain break is a pair of consecutive C*α* more than 4.0 Å apart. Counting instead the breaks a prediction introduces over the apo structure it was given, the median over the 150 test pairs is 8.0 for A2Holo against 101.5, 100.5 and 185.0 for FDBM, ABM and SBAlign, while A2Holo’s median gain in TM-score over that same input is the largest of the four (+0.111 against +0.070, +0.079 and −0.032, with +0.157 for the depositions). Pair by pair it introduces fewer new breaks than FDBM in 143 of 150 (*p* = 3.2×10*^−^*^36^, exact sign test, ties dropped), than ABM in 144 and than SBAlign in 149, and introduces none at all in 48 against 1, 2 and 0. Against the holo depositions themselves it introduces more in 101 of 150, tying in 48. These counts are for the released checkpoint at 10 steps and are C*α* only. No all-atom reconstruction is scored. In Figure A4, each row is the pair with the median A2Holo error at its difficulty level. None of the five is among the 6 pairs where FDBM’s best sample scores higher or the 30 where the untouched apo input does (25 by C*α* RMSD instead of TM-score).

To measure rotation consistency, we drew 100 Haar-uniform SO(3) rotations and applied each to the apo input. We then integrated the flow, rotated the prediction back into the original frame and compared it with the unrotated run, both as it stands and after superposing the two. The heaviest term of the objective, Equation 8, compares predicted with true displacement differences without superposing, so it constrains the orientation of the prediction. A global translation changes no term, and no reported metric sees rotation or translation, since all of them superpose. The predicted pose is therefore fixed up to a global translation, and the measurement separates the same way. Dropping the centroid offset takes the deviation from 0.51 to 0.078 Å, so nearly all of it is that translation. Superposing on top of that brings the back-rotated prediction within 0.075 Å of the unrotated run.

**Figure A4:**
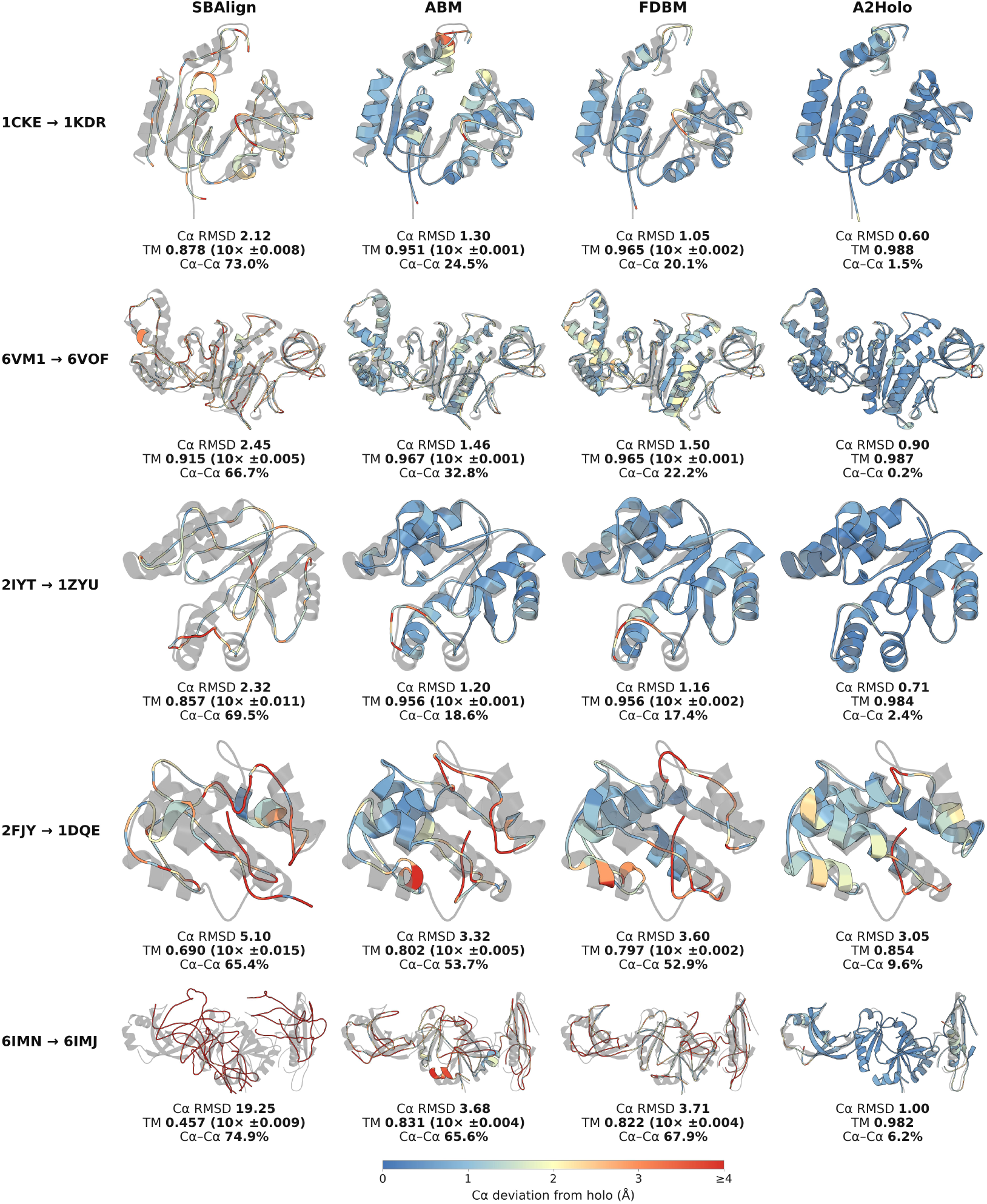
Predicted structures on five test pairs, one per difficulty level. Rows are test pairs, columns are methods. Each structure is coloured by its per-residue C*α* deviation from the holo deposition after Kabsch superposition on holo C*α*, on the single scale of the colourbar, and is drawn over the holo structure in translucent grey. The colouring, cap and grey are those of Figure 1. Under each panel: C*α* RMSD, TM-score, and the fraction of consecutive C*α*–C*α* distances more than 0.5 Å from 3.8 Å, over the bonds holo itself keeps. The three baselines keep the best of 10 samples, selected against the target. A2Holo integrates one deterministic trajectory (NFE 10, the released checkpoint) and has nothing to select among. For the baselines the TM-score is followed by ± half the range over their 10 samples. Rows were chosen before rendering, as the pair with the median A2Holo error at each of five difficulty levels, set by apo-to-holo C*α* RMSD over test pairs of 50–500 residues.

### A.9 Evaluation on a protein-disjoint split

In the D3PM split used in the main text, 123 of the 150 test pairs share a UniProt accession with a training or validation pair, and 112 share a PDB chain with a training pair. Therefore, we re-split all 1591 training, validation and test pairs of the D3PM split with the procedure of Zhang et al. (2023, Appendix C). Pairs are joined by single linkage when they share a UniProt accession or when their chains reach 0.7 in sequence similarity or TM-score. Test and validation each receive 20 whole clusters drawn at random with a fixed seed. Each then also receives the pairs of 10 single accessions taken from clusters whose other accessions stay in training. Test pairs are restricted to an apo–holo C*α* RMSD of 3–10 Å. We also apply this range to validation pairs, so that checkpoint selection sees the test distribution, and drop the 63 held-out pairs outside it rather than moving them to training. The result has 1425 training, 45 validation and 58 test pairs. We call it protein-disjoint because no test pair shares a UniProt accession or a PDB chain with a training or validation pair. One test pair shares a PDB entry with such a pair through a different chain.

A2Holo, SBAlign, ABM and FDBM are each retrained once (seed 1) on this split, A2Holo with the recipe of Appendix A.6. The baselines keep the checkpoint selected on the same validation pairs and draw 10 samples per test pair. They do not rank their own samples, so we keep one sample per pair by three rules (Table A1). We take the first sample, the sample with the fewest C*α*–C*α* bond violations or the sample closest to the holo structure. Only the last needs the holo structure. A2Holo runs one deterministic trajectory.

Per pair, A2Holo is more accurate than every baseline under every rule. It has the lower C*α* RMSD in 38–43 of the 58 test pairs (*p* ≤ 0.0045), the higher TM-score in 38–49 (*p* ≤ 8.0 × 10*^−^*^5^) and the higher lDDT in 48–57 (*p* ≤ 1.8 × 10*^−^*^7^), by two-sided Wilcoxon signed-rank tests. Its TM-score is also higher than the best of each baseline’s 10 samples ranked by TM-score, in 38–47 of the 58 pairs (*p* ≤ 2.2 × 10*^−^*^4^).

Table A2 compares backbone geometry, with each baseline keeping its geometry-best sample (Table A1), the choice most favourable to it. A2Holo moves the apo structure a median of 2.02 Å, against 2.15 Å for SBAlign, 2.20 Å for ABM and 2.62 Å for FDBM. A median of 3.4% of its consecutive C*α*–C*α* distances lie more than 0.5 Å from 3.8 Å, against 67.1% for SBAlign, 35.2% for ABM and 38.5% for FDBM.

**Table A1:**
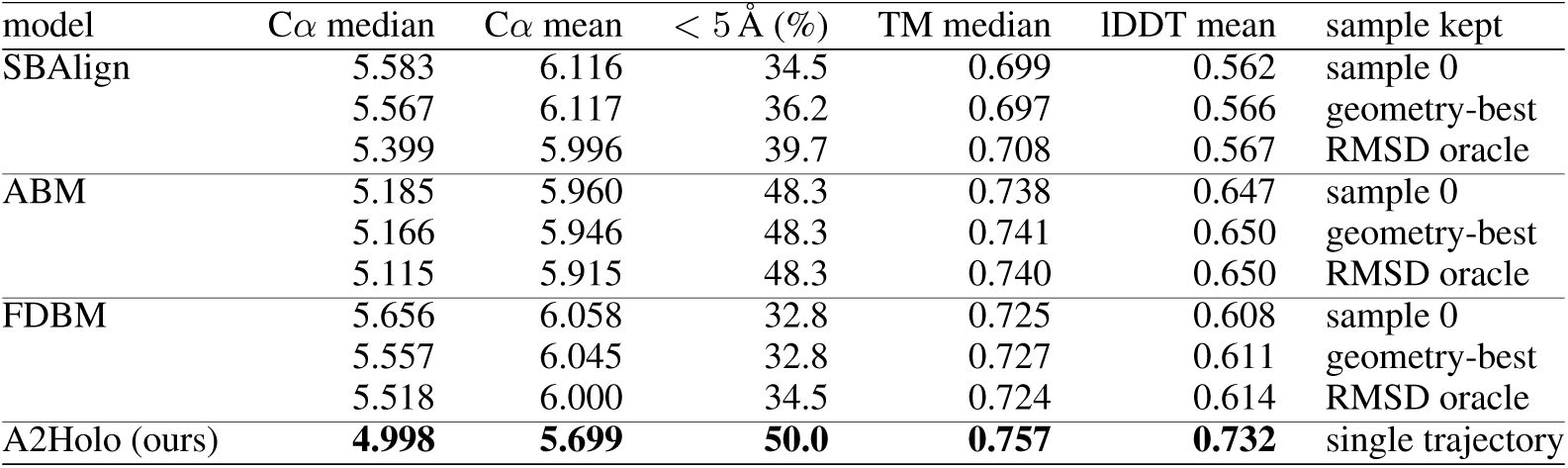
Accuracy on the 58 pairs of the protein-disjoint test set. Each baseline draws 10 samples per pair, and each of its rows keeps one by the rule named. Sample 0 is the first draw. Geometry-best is the sample with the fewest C*α*–C*α* bond violations and needs no holo structure. RMSD oracle is the sample closest to the holo structure and needs the answer. A2Holo runs one deterministic trajectory of 10 Euler steps. C*α* RMSD in Å. lDDT (Mariani et al., 2013) is computed on C*α* with a 15 Å inclusion radius and thresholds of 0.5, 1, 2 and 4 Å, and averaged over residues.

| model | Cα median | Cα mean | < 5 Å (%) | TM median | lDDT mean | sample kept |
|---|---|---|---|---|---|---|
| SBAlign | 5.583 | 6.116 | 34.5 | 0.699 | 0.562 | sample 0 |
|  | 5.567 | 6.117 | 36.2 | 0.697 | 0.566 | geometry-best |
|  | 5.399 | 5.996 | 39.7 | 0.708 | 0.567 | RMSD oracle |
| ABM | 5.185 | 5.960 | 48.3 | 0.738 | 0.647 | sample 0 |
|  | 5.166 | 5.946 | 48.3 | 0.741 | 0.650 | geometry-best |
|  | 5.115 | 5.915 | 48.3 | 0.740 | 0.650 | RMSD oracle |
| FDBM | 5.656 | 6.058 | 32.8 | 0.725 | 0.608 | sample 0 |
|  | 5.557 | 6.045 | 32.8 | 0.727 | 0.611 | geometry-best |
|  | 5.518 | 6.000 | 34.5 | 0.724 | 0.614 | RMSD oracle |
| A2Holo (ours) | **4.998** | **5.699** | **50.0** | **0.757** | **0.732** | single trajectory |

One further measurement on this split is reported in the conclusion, a rank correlation between attention concentration and off-target motion. Concentration is the measure of Appendix A.12, the attention a residue receives within the structure stream, ranked against its total distance change and partialled on its distance from the apo centroid, averaged over the double-stream blocks and taken at *t* = 0 with the apo structure as input. Off-target motion is the part of the predicted displacement that does not lie along the true one. Write **d**^t^ for the true displacement of a pair and **d**^p^ for the predicted one, each stacked over the residues and taken after Kabsch superposition on the apo coordinates. Then, with *r* = ⟨**d**^p^, **d**^t^⟩*/*⟨**d**^t^, **d**^t^⟩, off-target motion is ‖ **d**^p^ − *r* **d**^t^ ‖ */* ‖ **d**^t^ ‖. The correlation is Spearman over the 58 test pairs, Holm-adjusted within the 34 tests fixed before the run.

**Table A2:** Backbone geometry on the protein-disjoint test set, as medians over pairs of per-structure values (bond violations and angle outliers over the 57 pairs with valid bonds, clashes and displacement over all 58). A bond violation is a consecutive C*α*–C*α* distance more than 0.5 Å from 3.8 Å. A clash is a pair of C*α* closer than 3.4 Å and at least three residues apart. An angle outlier is a window of four consecutive C*α* whose virtual bond angle *θ* and dihedral *τ* fall in the sparsest (*θ, τ*) bins of the training holo structures, bins that together hold 0.5% of their windows. Displacement is the C*α* RMSD from the apo input. Each baseline keeps its sample with the fewest bond violations. SBAlign uses its released sampler, 10 Euler–Maruyama steps at a constant noise scale *g* = 1, and every step, the last included, adds Gaussian noise of total variance 1 Å^2^ per axis to each coordinate.

| model | bond violations (%) | clashes | angle outliers (%) | displacement (Å) |
|---|---|---|---|---|
| holo (deposited) | 0.0 | 0.0 | 0.4 | 5.22 |
| SBAlign | 67.1 | 22.5 | 39.7 | 2.15 |
| ABM | 35.2 | 14.5 | 12.3 | 2.20 |
| FDBM | 38.5 | 15.0 | 16.6 | 2.62 |
| A2Holo (ours) | **3.4** | **1.5** | **1.5** | 2.02 |

**Table A3:**
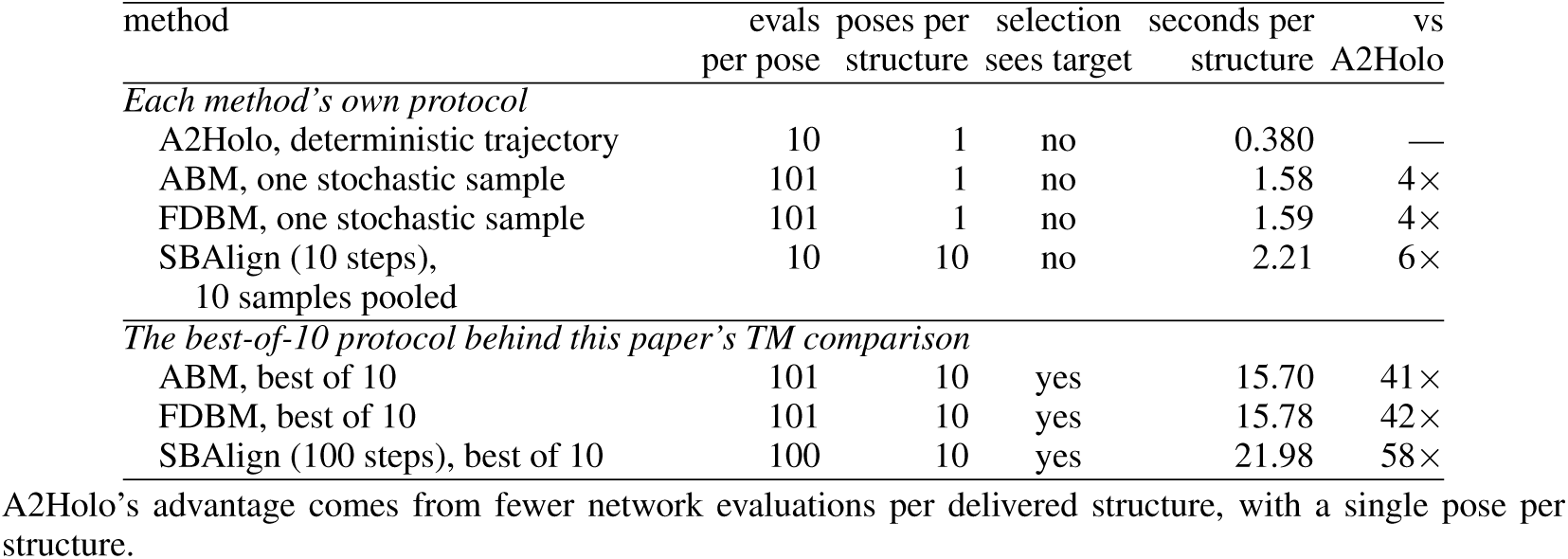
Inference cost of the one structure each method actually delivers, not of a matched step count. Evals counts network forward passes per pose, and selection marks picking the pose closest to the holo target. Per-pair wall clock over the 150 test pairs, median over pairs, one NVIDIA RTX A5000, one session. It times the inference call only, with the protein language-model embedding computed outside the timed region. A2Holo’s trajectory is deterministic, so its single draw is also what the TM comparison used.

| method | evals | poses per | selection | seconds per | vs |
|---|---|---|---|---|---|
|  | per pose | structure | sees target | structure | A2Holo |
| *Each method’s own protocol* |  |  |  |  |  |
| A2Holo, deterministic trajectory | 10 | 1 | no | 0.380 | — |
| ABM, one stochastic sample | 101 | 1 | no | 1.58 | 4× |
| FDBM, one stochastic sample | 101 | 1 | no | 1.59 | 4× |
| SBAlign (10 steps), 10 samples pooled | 10 | 10 | no | 2.21 | 6× |
| *The best-of-10 protocol behind this paper’s TM comparison* |  |  |  |  |  |
| ABM, best of 10 | 101 | 10 | yes | 15.70 | 41× |
| FDBM, best of 10 | 101 | 10 | yes | 15.78 | 42× |
| SBAlign (100 steps), best of 10 | 100 | 10 | yes | 21.98 | 58× |

A2Holo’s advantage comes from fewer network evaluations per delivered structure, with a single pose per structure.

### A.10 Inference cost

Table A3 times the inference call alone for every method, on the same 150 test pairs and the same NVIDIA RTX A5000. For A2Holo the Ankh3-large embedding of the sequence adds a median of 77 ms per chain on the same GPU model, so its full cost is 0.456 s per pair, 35× below FDBM’s best of 10. No baseline uses a language model, and every row excludes the same offline preprocessing.

Every stochastic baseline appears in both halves of the table. SBAlign’s upper row follows its published protocol, which pools 10 samples and takes their median, as in Table 1. Its lower row gives it the best-of-10 selection used in the TM comparison, at 100 Euler steps instead of 10.

### A.11 Statistics

Summary statistics are medians over the 150 test pairs. That matches the convention used in earlier reports on this benchmark and limits the influence of the few pairs with very large binding-induced motion. Unless stated otherwise, paired comparisons between methods are made on per-pair values with the two-sided Wilcoxon signed-rank test (Wilcoxon, 1945), using the same pairs for both methods. No pair was excluded from the benchmark evaluation. The measurements of Section 4 instead use the 133 test pairs under a length cutoff, which bounds the size of the joint attention map. Appendices A.9 and A.15 use the 58 test pairs of the protein-disjoint split.

Interval estimates in Section 4 are percentile bootstraps over pairs. The pair set is resampled 2000 times with replacement, the statistic is recomputed as the mean over each resample, and the interval is taken at the 2.5th and 97.5th percentiles under a fixed seed. They are descriptive and carry no test.

Where two quantities measured on the same training are compared, each training contributes one event, *Z_m_* = ⃦[the first exceeds the second in training *m*], and the count *_m_ Z_m_*is tested with a two-sided sign test.

### A.12 The attention quantities, defined

Section 4.2 defines the emitted and received attention of a residue and the identity that links a residue’s two emitted vectors.

The sum 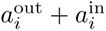 of the structure-to-structure quadrant is the natural per-residue summary, since both ends index the same residue, but it does not separate the two directions. An unrestricted row sum is identically one, because rows are normalised over all 2*N* keys, so the emitted sum is taken inside one quadrant.

Equation 4 holds under two restrictions. It constrains the emitted, or row, vectors only. Column sums are unconstrained and no analogous identity exists for received attention. The index must also be fixed to one stream. For a structure-stream query the *j* ≤ *N* term is the cross-stream one, and for a sequence-stream query the roles swap. Because the two emitted vectors sum to 1*/N*, the top 15% of the cross-stream one is the bottom 15% of the intra-stream one. Its overlap with the moving residues therefore differs from the intra-stream overlap in size while carrying no additional information.

Rank correlations *ρ* in this paper are Spearman’s, with average ranks for ties. Because *x* ’→ 1*/N* −*x* reverses every rank and maps tied values to tied values, *ρ*_cross_ = − *ρ*_intra_ stays exact with ties, and it would hold equally for Pearson’s coefficient. Equation 4 is exact in exact arithmetic, and it holds in the measured data. Across all 108 model × layer cells of the structure stream the two emitted Spearman correlations sum to at most 1.000 × 10*^−^*^5^ in absolute value, the precision at which these tables are stored. The sequence stream matches it, and it is the structure stream the paper relies on.

The entrywise correlation of Section 4.3 is Pearson’s *r*. The conditional analysis of Section 4.3 holds fixed *g_i_*, the distance of residue *i*’s C*α* from the apo C*α* centroid. Its partial correlation is the first-order partial of Spearman coefficients,

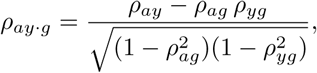

where **a** is the attention vector and *y_i_* =∑*_j_* |Δ*D_ij_*|, computed per pair and block and averaged like every other quantity. The residual overlap drops the top 15% of residues by *g_i_* and compares, among the rest, the top 15% by attention with the top 15% by *y_i_*, against the same chance level *f/*(2 − *f*). Appendix A.9 uses this partial correlation again on the protein-disjoint split, kept per pair and taken at *t* = 0.

Section 4.4 reads each cross quadrant at its structure-residue end. Each cross quadrant has two ends, and exactly one of them is indexed by a structure-stream residue. In *A*^(*ℓ*)^[*N* :, :*N*] the rows are structure-stream residues, which read from the sequence stream, and the columns are sequence tokens. In *A*^(*ℓ*)^[:*N, N* :] the rows are sequence tokens and the columns are structure-stream residues, which the sequence stream reads. Figure A5 draws these quadrants on five pairs, across blocks. Figure 5 draws four of these quantities on one pair, at five flow times.

**Figure A5:**
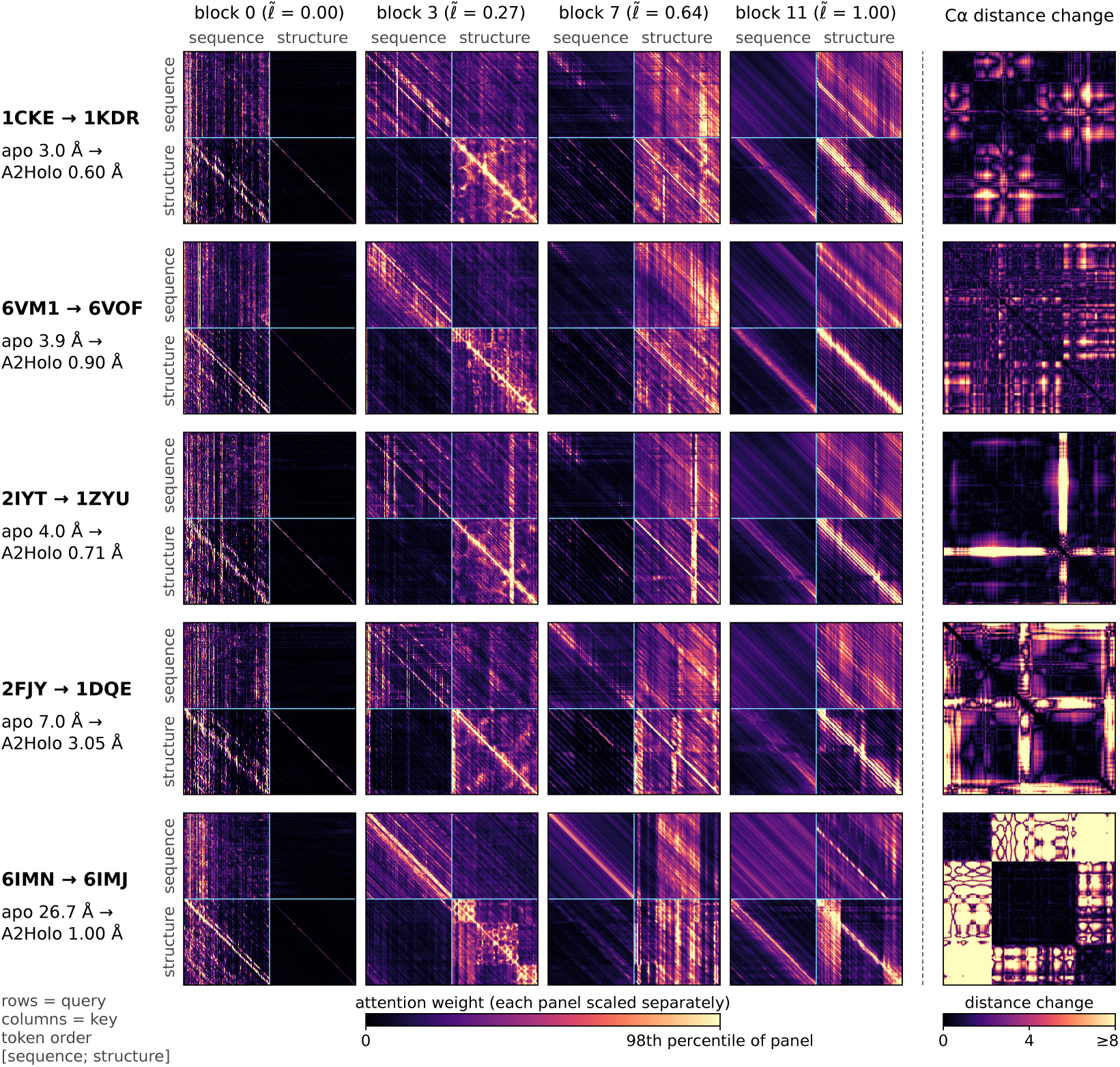
The object this paper measures. Head-averaged joint attention of the released model at *t* = 0.5, on the five test pairs of Figure A4, at four double-stream blocks evenly spaced through the stack. Blocks are numbered from 0, with normalised depth *ℓ̃* = *ℓ/*(*L* − 1). In each map rows are queries and columns are keys, in the model’s token order [sequence; structure], so the top-left quadrant is sequence-to-sequence and the bottom-right is structure-to-structure. The last column is each pair’s C*α* distance change |*D*_holo_ − *D*_apo_|, capped at 8 Å. Each map is scaled to its own 98th percentile, so the maps show where attention sits and not how much of it there is. In the first block the structure queries read almost only the sequence stream. Later blocks build a sharp structure-to-structure diagonal and bright structure columns, the residues that are read most (received attention, Section 4.2). The maps are illustrations, and claims are made over the 9 trainings and 133 test pairs (Figure 6).

**Figure A6:**
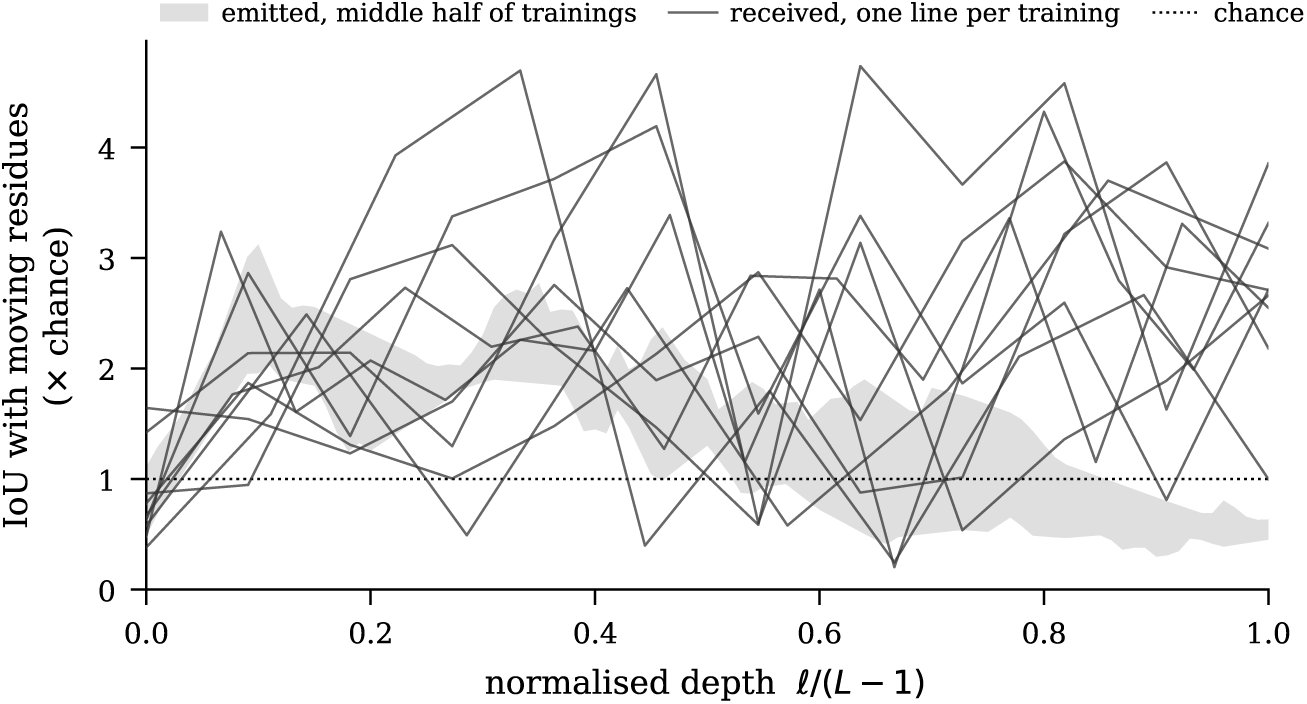
Depth profiles behind the depth means of Figure 6. Overlap between structure-to-structure attention and the moving residues, as a multiple of chance, against normalised depth *ℓ/*(*L* − 1), over 133 test pairs at *t* = 0.5. Each dark line is one of the 9 trainings, for received attention. The grey band spans the middle half of the trainings for emitted attention. The profiles cross and peak at different depths, which is why the text quotes depth means (Section 4.1).

**Table A4:** Every training of Section 4, evaluated as Table 1 evaluates the released one, deterministically with NFE 10. The upper block varies only the random seed at the released depth of 12+12 blocks. The lower block varies the depth of both stacks, at seed 1. The released checkpoint is seed 1.

|  | Cα RMSD (Å) |  |  |  |
|---|---|---|---|---|
| Training | Median | Mean | <2 | <5 |
| **seed 1 (released)** | **0.809** | **2.357** | **66.7%** | **88.0%** |
| seed 2 | 0.761 | 2.401 | 64.7% | 87.3% |
| seed 3 | 0.836 | 2.491 | 65.3% | 86.0% |
| seed 4 | 0.777 | 2.377 | 66.0% | 88.0% |
| seed 5 | 0.774 | 2.419 | 66.0% | 86.0% |
| depth 8 | 0.747 | 2.458 | 64.7% | 87.3% |
| depth 10 | 0.826 | 2.381 | 64.7% | 88.7% |
| depth 14 | 0.835 | 2.374 | 64.7% | 87.3% |
| depth 16 | 0.781 | 2.384 | 64.7% | 88.7% |

The overlap with the moving residues is a depth average and is never read at an argmax over 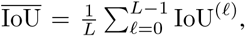, where IoU^(*ℓ*)^ = |*S*^(*ℓ*)^ ∩ *M* |*/*|*S*^(*ℓ*)^ ∪ *M* | is the intersection over union of two residue sets of equal size [*f N* |, averaged over pairs. *S*^(*ℓ*)^ holds the residues ranked highest by an attention vector at block *ℓ*, a row or a column mean of one quadrant, and *M* those ranked highest by ∑*_j_* |Δ*D_ij_*|. For two top-*f* sets of equal size the chance level is *f/*(2 − *f*), which is 0.0811 at *f* = 0.15. Measured on random sets, the overlap is 1.06 times this level, because *f/*(2 − *f*) is the expected intersection over the expected union rather than the mean of the ratio. Figure A6 draws the per-training depth profiles behind these depth means. They cross and peak at different depths.

### A.13 Retraining variability

The recipe reported here was trained 5 times under one configuration, differing only in the random seed, and the test median C*α* RMSD spans 0.761–0.836 Å, a spread of 0.075 Å (Table A4).

The 4 depth variants (Table A4, lower block) are single trainings at seed 1. The depth variants are among the 9 trainings so that a block index does not mean the same depth in every model, and Section 4 makes no claim about depth as a variable.

### A.14 Removing the sequence stream

The sequence stream was removed and the model trained from scratch under the recipe and stopping rule of Appendix A.6 (Table A5). Removing the stream costs 0.195 Å of test median C*α* RMSD, 24%, more than the 0.075 Å spread across retrainings of the released configuration (Appendix A.13). Without the sequence stream there is nothing for a double-stream block to join, so this model keeps only the single-stream blocks. Therefore, the 0.195 Å also includes the loss of the double-stream stack.

**Table A5:** Removing the sequence stream. Test C*α* and C*β* median RMSD over the 150 test pairs at 10 deterministic Euler steps.

| model | Cα median | Cβ median | Δ Cα |
|---|---|---|---|
| A2Holo (ours) | 0.809 | 0.992 | — |
| w/o sequence stream | 1.004 | 1.461 | 0.195 |

### A.15 Sequence information, fusion and model size

The 0+12 architecture of Appendix A.14 has 113.7M parameters, against 368.8M for A2Holo. To separate sequence information, fusion and model size, we train seven models on the protein-disjoint split of Appendix A.9 (Table A6), among them a parameter-matched model (367.8M) without language-model inputs. Four differ in model size and in whether they receive the language-model embedding. Two keep the Ankh3 sequence stream, one with the released 12+12 blocks and one with 4+4 blocks. Two have no language-model embedding. One has the 0+12 single-stream architecture of Appendix A.14. The other keeps the 12+12 dual-stream architecture, but its second stream receives a copy of the structure input in place of the embedding and there is no global token. It differs from A2Holo only in lacking these two Ankh3 inputs. Residue identity remains in the node features of every model. A fifth model tests the fusion. It keeps A2Holo’s architecture and size but replaces the joint attention of each double-stream block with cross-attention, in which each stream attends only to the other. Two more test the language model and replace Ankh3 with ESM-2 or ESM-C in the released 12+12 architecture. Five of the seven models have about the same size, 367.8 to 368.8M parameters. All seven use the recipe of Appendix A.6, one training each (seed 1).

With the sequence stream, the larger model is more accurate. Going from 4+4 to 12+12 lowers the C*α* RMSD in 37 of the 58 test pairs (*p* = 0.0037) and the C*β* RMSD in 37 (*p* = 0.0026). Without it, the larger model is less accurate. Going from 0+12 to the 12+12 structure copy raises the C*α* RMSD in 36 pairs (*p* = 8.4 × 10*^−^*^4^). The two effects differ in 38 pairs (*p* = 3.1 × 10*^−^*^4^, two-sided Wilcoxon signed-rank tests throughout). Therefore, the larger dual-stream model pays off only when its second stream carries the language-model embedding. The two models without the embedding move the apo structure less than the five with it (median displacement 0.71–0.73 Å against 1.12–2.02 Å). At 12+12, A2Holo moves it further than the structure copy in 54 of the 58 pairs (*p* = 2.7 × 10*^−^*^10^). Therefore, every language-model embedding makes the model move the apo structure. A2Holo has the lowest median C*α* and C*β* RMSD of the seven models (Table A6).

### A.16 The first double-stream block and block removal

At the first double-stream block, where the two streams first meet, a structure-stream residue pays most of its attention to the sequence stream in all 9 trainings. No later block does so in every training. Attention is also most concentrated at that first block in 6 of the 9 trainings, by row entropy, the Shannon entropy of each attention row divided by log 2*N* and averaged over rows.

Section 4.5 also removes one block’s residual update at a time, and Figure A7 draws what that costs, <u>o</u>nce pe<u>r</u> training. With *δ_ℓ_*the rise in C*α* RMSD when block *ℓ*’s update is removed, the ratio Γ = *δ*_double_*/δ*_single_ of the mean costs runs from 0.8 to 10.3, over per-block costs whose ranges overlap. The two removals differ in kind, since a double-stream block keeps its sequence stream while a single-stream one loses its whole concatenated update. At the boundary, where depth position is nearly fixed, the double block costs more in 2 of 9 trainings (*p* = 0.18), so this test does not resolve an ordering.

**Table A6:** Accuracy of the seven models on the 58 test pairs of the protein-disjoint split (Appendix A.9). The first row is A2Holo, the same model as in Table A1. One training per model (seed 1), best validation checkpoint, 10 Euler steps. Blocks are double-stream + single-stream, and attention is that of the double-stream blocks. RMSDs in Å. Displacement is the median C*α* RMSD of the prediction from the apo input.

| second stream | blocks | attention | Cα median | Cβ median | parameters (M) | displacement (Å) |
|---|---|---|---|---|---|---|
| Ankh3 (A2Holo) | 12+12 | joint | **4.998** | **5.128** | 368.8 | 2.02 |
| *size and second stream* |  |  |  |  |  |  |
| Ankh3 | 4+4 | joint | 5.362 | 5.611 | 127.9 | 1.12 |
| none | 0+12 | — | 5.235 | 5.428 | 113.7 | 0.71 |
| structure copy | 12+12 | joint | 5.283 | 5.596 | 367.8 | 0.73 |
| *fusion* |  |  |  |  |  |  |
| Ankh3 | 12+12 | cross | 5.239 | 5.541 | 368.8 | 1.85 |
| *protein language model* |  |  |  |  |  |  |
| ESM-2 | 12+12 | joint | 5.338 | 5.732 | 368.4 | 1.74 |
| ESM-C | 12+12 | joint | 5.365 | 5.799 | 367.9 | 1.97 |

**Figure A7:**
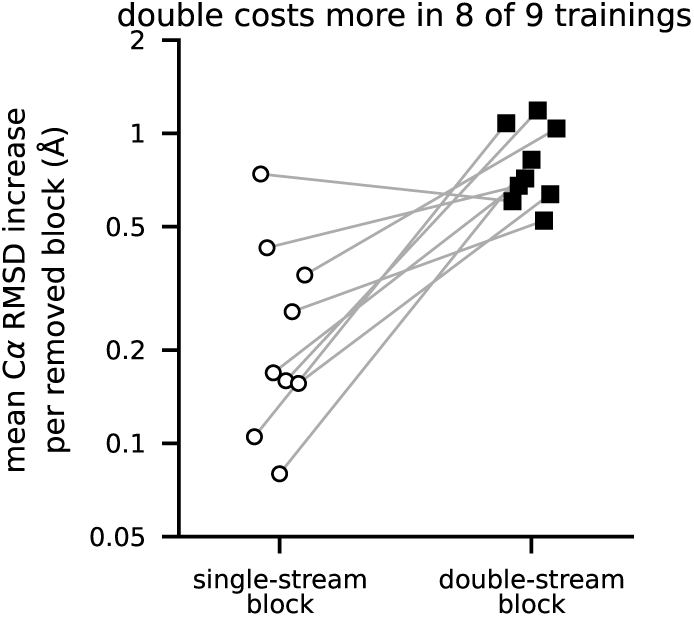
Removing a double-stream block costs more than removing a single-stream one in 8 of 9 trainings. The cost is the rise in C*α* RMSD when one block’s residual update is taken out, averaged over the blocks of that type and drawn on a log scale. A line joins the two block types of one training.

#### Quadrant damping

In one double-stream block at a time, the two cross-stream quadrants of the joint attention map are multiplied by *α* and every row is renormalised to sum to one, so attention is re-routed rather than removed. The control scales the two intra-stream quadrants instead, which touches the same number of entries. Parameters and tokens are unchanged. The cost is the rise in C*α* RMSD, against holo, of the one-step prediction **x**_apo_ + **v***_θ_*(**x***_t_, t*) at *t* = 0.5 over the 133 test pairs, averaged over pairs and then over the double-stream blocks of each training. *α* = 0 zeroes a quadrant and *α* = 0.5 halves it.

### A.17 Why the probes take the form they do

#### Global-frame displacement is not a well-posed probe target

Regressing Δ**x***_i_* itself from a hidden state is ill-posed for a reason unrelated to the model. Each apo–holo pair is deposited in its own arbitrary crystallographic frame. The *direction* of Δ**x***_i_* therefore carries no meaning shared across proteins, and a single linear map has no frame in which to generalise. Section 4.5 uses |Δ**x***_i_*| and the total distance change ∑*_j_* |Δ*D_ij_*| instead, both invariant to the frame (Figure A8).

Invariant targets carry a different risk. They could recover a flexibility prior the sequence already fixes. The denominator of Equation 5, a probe of the same family fitted on the frozen language-model embedding alone, controls for it by construction. On its own, that probe reaches *R*^2^ = 0.089 for |Δ**x***_i_*| and 0.203 for total distance change. Each probe is an affine ridge map with an unpenalised intercept, fitted in closed form on unstandardised features. Its input **h**^(*ℓ*)^ is the state of residue *i*’s structure token. In the single-stream stack there is no separate structure stream, and **h**^(*ℓ*)^ is then the continuation of that token in the merged sequence. The 133 test pairs are assigned to five folds by pair, so every residue of a pair stays in one fold. Pairs that share a protein can still fall in different folds. Probes share this architecture and these folds across all blocks, and the sequence-only denominator is fitted by the same procedure on the same folds. The backbone is frozen throughout. The ridge penalty is chosen per probe, on the same held-out folds the probe is then scored on. Choosing it instead by an inner cross-validation over the other folds, for the numerator and the sequence-only denominator alike, leaves every value of the 9 trainings that Section 4.5 prints unchanged. The expectation in Equation 5 is the squared error per residue, pooled over all held-out residues rather than averaged per protein, so longer proteins weigh more.

**Figure A8:**
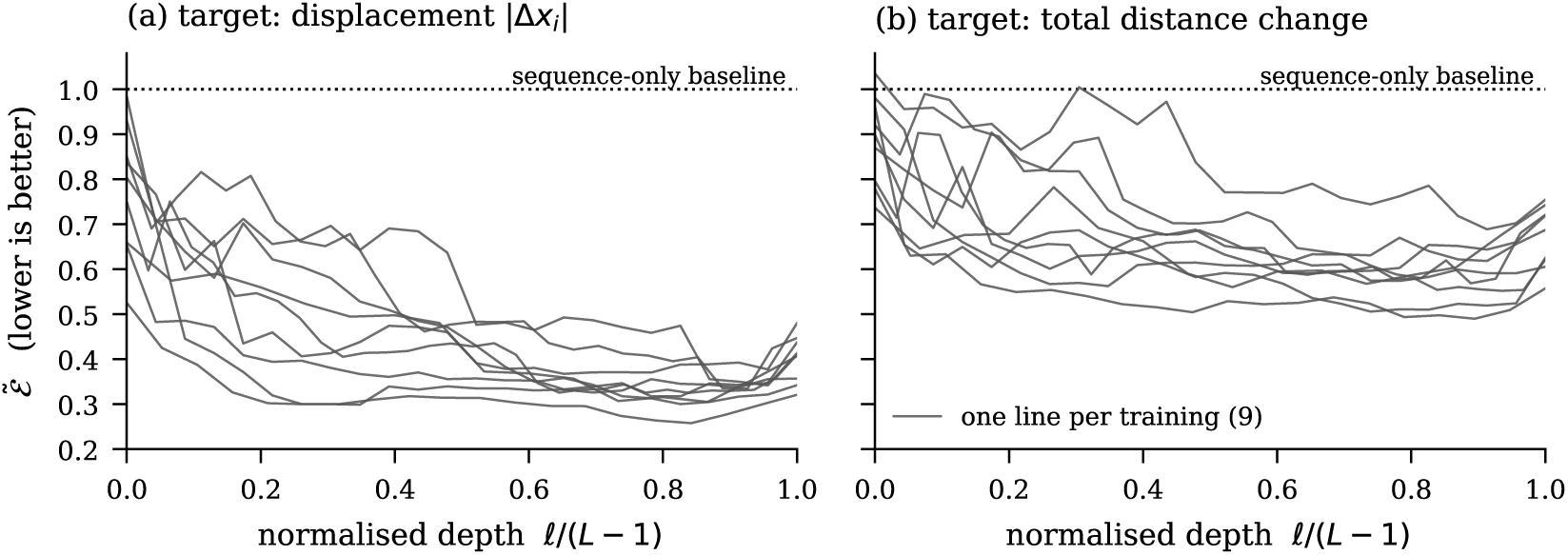
What each depth adds over what the sequence already determines. Relative decodability Ẽ (Equation 5) is plotted for the two rotation-invariant targets. The horizontal axis is normalised depth *ℓ/*(*L* − 1) over both stacks, since the 9 trainings mix depths. Each line is one of the 9 independently trained models. Both panels share one vertical scale. The dotted line at 1 is the sequence-only probe, so lower is better. Averaged over depth, Ẽ runs 0.320–0.569 in (a) and 0.549–0.845 in (b), below one in all 9 trainings for both (Section 4.5).

#### Random-initialisation control

At each of the 5 depths, 3 untrained networks of that architecture are probed exactly as above, with the same targets, folds and sequence-only denominator. Their depth means are 1.071–1.204 for displacement magnitude and 1.034–1.187 for total distance change, against 0.320–0.569 and 0.549–0.845 for the trainings, so an untrained stack is less informative than the sequence alone. At *t* = 0 the untrained bands are 1.005–1.229 and 0.964–1.181, and every training again sits below the untrained mean at its own depth (Table A7).

#### The minimum over depth

Each model’s best block reaches Ẽ = 0.258–0.367 for displacement magnitude and 0.490–0.688 for total distance change. The block that attains it is chosen per model, so the value is a magnitude and carries no test.

### A.18 Every Section 4 quantity at *t* = 0

Section 4 measures at *t* = 0.5, on the interpolant between the apo and holo structures. The deployed model starts from the apo structure alone, so every measurement was repeated at *t* = 0 on the same 9 trainings, with the analysis unchanged.

Table A7 gives the result behind each headline claim of Section 4 at both flow times. Five counts over the trainings move at *t* = 0, each by one training. The cross-stream read of displacement and the structure-to-sequence end comparison each lose one training. Section 4.4 already leaves both unresolved. The first block’s Ẽ for total distance change falls below one in one more training, and at the boundary of Appendix A.16 the double block costs more in one more training, still unresolved. The cross-stream correlation against untrained networks of Appendix A.19 also loses one training. Every other count over the trainings in Table A7, and its *p*, is unchanged. The last row, which counts random initialisations, rises from 2 to 8 of 24, while all 9 trainings keep that ordering. The ranges shift at their lower ends, most for the depth of the lowest Ẽ, a location Section 4 already reports as not reproducing.

**Table A7:** The result behind each headline claim of Section 4, at *t* = 0.5 and at *t* = 0. At *t* = 0 the model is given the apo structure alone, the deployed operating point, and the *t* = 0.5 column repeats the values printed in Section 4. Counts are *k/N* over trainings with the sign-test *p* where Section 4 prints one. The last row counts random initialisations and carries no test. Ranges are absolute values over trainings.

| claim | *t* = 0.5 | *t* = 0 |
|---|---|---|
| Structure stream: received > emitted | 9/9, *p* = 0.0039 | 9/9, *p* = 0.0039 |
| above own-depth random-init mean | 9/9, *p* = 0.0039 | 9/9, *p* = 0.0039 |
| Partial *ρ*, centroid fixed, > random-init mean | 9/9, *p* = 0.0039 | 9/9, *p* = 0.0039 |
| range of that partial *ρ* | 0.046–0.145 | 0.037–0.146 |
| Sequence→structure > sequence self | 9/9, *p* = 0.0039 | 9/9, *p* = 0.0039 |
| $\tilde{\mathcal{E}}$ < 1, displacement; distance difference | 9/9; 9/9, *p* = 0.0039 | 9/9; 9/9, *p* = 0.0039 |
| $\tilde{\mathcal{E}}$ below random-init mean, two targets | 9/9; 9/9, *p* = 0.0039 | 9/9; 9/9, *p* = 0.0039 |
| Cross-stream damping > intra-stream | 8/9, *p* = 0.039 | 8/9, *p* = 0.039 |
| Double-block removal > single-block | 8/9 | 8/9 |
| Attention vs. distance change, *ρ* > 0 | 8/9, *p* = 0.039 | 8/9, *p* = 0.039 |
| Depth $\tilde{\ell}$ of lowest $\tilde{\mathcal{E}}$, two targets | 0.348–0.957; 0.348–0.935 | 0.261–0.957; 0.217–0.913 |
| Cross-stream read vs. \|Δ**x**<sub>*i*</sub>\|, *ρ* > 0 | 6/9, *p* = 0.51 | 5/9, *p* = 1.00 |
| Structure-indexed received > sequence-indexed | 9/9, *p* = 0.0039 | 9/9, *p* = 0.0039 |
| same ordering at random initialisation | 2/24 | 8/24 |

The geometric baselines take no model input, so one computation serves both flow times. They select residues from the apo structure alone and are scored against the same moving residues.

### A.19 The rest of the limitations

#### Scope of Section 4

The attention results of Section 4 are associations between attention statistics and apo–holo targets. The interventions (quadrant damping, block removal and removing the sequence stream) show what the prediction depends on, not that attention is what places the change. The conditional analysis of Section 4.3 controls one geometric variable, distance from the apo centroid. The cross-stream correlation with |Δ**x***_i_*| (Section 4.4) exceeds the mean of untrained networks of the same depth in 6 of 9 trainings (*p* = 0.51), which this test cannot resolve. The correlation with how much a residue is read by its own stream exceeds it in 8 of 9 (*p* = 0.039). Each measurement is a snapshot at one flow time rather than an integral along the path, and the sign tests of Section 4 share the same 9 trainings without correction for multiplicity.

#### Model output

Only C*α* and C*β* are predicted, so a full-atom model needs an external reconstruction step. Here the C*α*-to-all-atom network cg2all (Heo & Feig, 2024) is used only to draw Figures 1, 3, 5 and A4 and is not evaluated. Scoring full-atom models and using the predicted C*α* trace as a template for a structure predictor such as Boltz-2 (Passaro et al., 2025) are left to future work.

#### Single state

A deterministic pose cannot represent an ensemble or a second bound state. The model sees one chain and no ligand, so it opens a pocket without knowing what will occupy it.

#### Rotation consistency

Rotation consistency is measured, not guaranteed. Over 100 rotations of the released model the predicted shape moves by 0.075 Å and its C*α* RMSD to holo by 0.005 Å. The test median stays between 0.801 and 0.815 Å, and 1 of 150 pairs moves by more than half an ångström under at least one of them. The model gets this consistency from rotation augmentation rather than an equivariant architecture.

